# Individualized network topography in pre-adolescent children and adults using naturalistic precision fMRI

**DOI:** 10.64898/2026.03.05.709899

**Authors:** Shefali Rai, Kate J. Godfrey, Kirk Graff, Ryann Tansey, Daria Merrikh, Shelly Yin, Matthew Feigelis, Damion V. Demeter, Tamara Vanderwal, Deanna J. Greene, Signe Bray

**Affiliations:** Child and Adolescent Imaging Research Program, University of Calgary, Calgary, AB, Canada; Alberta Children’s Hospital Research Institute, University of Calgary, Calgary, AB, Canada; Hotchkiss Brain Institute, University of Calgary, Calgary, AB, Canada; Department of Neuroscience, University of Calgary, Calgary, AB, Canada; Department of Radiology, University of Calgary, Calgary, AB, Canada; Department of Community Health Sciences, University of Calgary, Calgary, AB, Canada; Department of Cognitive Science, University of California San Diego, La Jolla, CA, USA; Department of Psychiatry, Faculty of Medicine, University of British Columbia, Vancouver, Canada

**Keywords:** Precision fMRI, Functional Brain Networks, Functional Connectivity, Naturalistic fMRI, Developmental Neuroimaging

## Abstract

Group-averaged network definitions, commonly used in developmental functional connectivity research, limit our understanding of how network topography may change with age and can lead to inaccurate estimates of age effects on intra- and inter-network functional connectivity. Here, we collected a precision fMRI dataset from 24 parent-child pairs (children 6 to 8 years, 13 females; adults 33 to 47 years, 12 females) during passive video viewing and derived individual template-matched functional network maps from 60 minutes of motion-censored data per participant. Overall, large-scale network architecture was broadly shared between children and adults, with no age-effects observed for network-level surface area, few age differences in network boundaries and generally lower assignment confidence in children. Considering inter-individual similarity in network topography, association networks showed stronger within-family similarity and no overall effect of age on similarity of networks between pairs of individuals. We further asked how individualized network definitions might impact estimates of age effects on dense functional connectivity. While all approaches pointed to greater within-network functional connectivity in adults, we found that individualized approaches had larger age effects and lower sensitivity to head motion. Together, our results suggest that, relative to adults, pre-adolescent children show reduced network assignment confidence and weaker within-network connectivity, but limited differences in network borders and size, and underscore the value of individualized mapping for increasing sensitivity to age effects.

## Introduction

Describing how functional brain networks change across childhood can provide insights into cognitive and behavioral maturation and establish a foundation for clinical investigations (Fair et al., 2009; Grayson & Fair, 2017; Satterthwaite et al., 2018). Studying developmental effects on functional connectivity has been challenging, however, as pediatric scans tend to have high head motion (Engelhardt et al., 2017; Frew et al., 2022; Satterthwaite et al., 2012), obtaining longer scans needed to optimize test-retest reliability presents challenges to compliance (Barkovich et al., 2019; Raschle et al., 2012), and the network templates used to define FC and interpret within- and between-network differences have largely been derived from groups of adults (Power et al., 2011; Thomas Yeo et al., 2011). Precision fMRI (Gordon, Laumann, Gilmore, et al., 2017; Gratton & Braga, 2026) a technique in which relatively long fMRI timeseries are collected per individual in order to achieve reliable mapping of networks and connectivity estimates, can help to circumvent these issues and ultimately improve our understanding of how functional connectivity strength and topography associates with age. Here, we use precision fMRI in a sample of pre-adolescent children and adults to investigate age effects on individually defined network topography and dense functional connectivity.

Functional network topography refers to the spatial location of functional networks (Demeter & Greene, 2025; Gordon, Laumann, Gilmore, et al., 2017; Lynch et al., 2020). While canonical network topography is broadly similar across individuals (Damoiseaux et al., 2006), precision functional mapping in adults has shown profound and reliable inter-individual variations (Dworetsky et al., 2024; Gordon, Laumann, Gilmore, et al., 2017; Gratton, Kraus, et al., 2020; Kwon et al., 2025; Seitzman et al., 2019), with greater inter-variability in association relative to sensory networks (Braga & Buckner, 2017; Kong et al., 2019). Emerging literature is using precision approaches to describe topography in children (Keller et al., 2023), including in infants (Labonte et al., 2024; Moore et al., 2024). However, while some work has combined across datasets to consider age effects on network boundaries across adolescence (Sun et al., 2025; Tooley, Bassett, et al., 2022; Tooley, Park, et al., 2022), data are limited on whether and how networks defined precisely in individual adults and children may differ (Demeter et al., 2025), a gap we aim to fill here.

To address the challenge of compliance in acquiring low-motion fMRI across multiple sessions in children, we chose passive viewing over task-free rest during fMRI acquisition. Passive viewing improves compliance and reduces head motion in pediatric populations (Greene et al., 2018; Vanderwal et al., 2015), and yields reliable individualized connectomes (Jiahui et al., 2023; O’Connor et al., 2017; Rai et al., 2025; Shearer et al., 2025; Wang et al., 2017). Further, naturalistic paradigms maintain intrinsic FC patterns and network architecture while offering greater ecological validity (Bray et al., 2015; Finn et al., 2017; Vanderwal et al., 2017) and can increase sensitivity in FC-behavior associations (Finn & Bandettini, 2021). Here, we asked whether network topography defined during passive viewing differs between adults and pre-adolescent children.

Specifically, we compared individualized network organization between pre-adolescent children (6.56 – 8.92 years) and parents recruited as adult controls, with familial relationships considered in analyses. We applied template-matching to derive individualized network maps and assessed age-related differences in topography, network assignment confidence, network size, and spatial location. We tested whether topography of parent-child pairs were more similar than unrelated pairs, given prior evidence of heritability in FC and network topography (Anderson et al., 2021; Sinclair et al., 2015), and whether adults or children had more individually unique network topography. Finally, we examined whether individualized network definitions improved sensitivity to age-related differences in FC.

## Methods

### Participants

The PreciseKIDS study recruited 25 parent-child pairs, comprising 5 female-female, 5 male-male, 7 female-male, and 8 male-female dyads. At the first session, children were aged between 6.56 and 8.92 years (mean = 7.88 years, s.d. = 0.69 years; 13F) and one of their parents aged between 33.75 and 47.13 years (mean = 41.39 years, s.d. = 3.63 years; 12F). One parent-child pair was excluded from this study due to excessive motion (< 40 minutes of usable data for the child), resulting in a final sample of 24 pairs. Two families contributed data from two parents and two children. Participants were recruited through community advertisements and a database of participants from previous studies who provided consent to contact for future research. Exclusion criteria were self-reporting a diagnosis of neurological, mental health condition, brain injury or chronic health condition. Children gave assent, and parents provided written informed consent for their own and their child’s participation. All procedures were approved by the University of Calgary Conjoint Health Research Ethics Board (REB19-1973). As done previously (Rai et al., 2025), we classified motion groups based on a threshold of 3863 volumes, resulting in a low-motion adult group (LMA) (n = 22, 11 M, 11 F; aged 33.75 – 47.13 years; mean FD: 0.048 – 0.163), a low-motion child group (LMC) (n = 10, 4 M, 6 F; aged 6.56 – 8.92 years; mean FD: 0.050 – 0.270), and a high-motion child group (HMC) (n = 14, 8 M, 6 F; aged 7.03 – 8.90 years; mean FD: 0.122 – 0.939) for analyses, where appropriate.

### MRI data collection

Parent-child pairs each completed four MRI sessions at the Alberta Children’s Hospital in Calgary, Canada, approximately 3 to 4 weeks apart (mean = 25.9 days between visits). Scans were conducted on a 3T GE MR750w system (Waukesha, WI) with a 32-channel head coil. EEG and cognitive assessments were also collected as part of the study but not reported here. Session order was pseudo-randomized, i.e., for half of the families, parents underwent EEG followed by MRI while their children completed MRI first, then EEG; the order was reversed for the other half of the families. Each MRI session included a T1-weighted 3D BRAVO sequence (0.8 mm isotropic voxels, TR = 6.764 ms, TE = 2.908 ms, FA = 10°). Functional imaging data were acquired in six runs per session using multi-echo, multiband T2*-weighted gradient-echo echo-planar sequence (ME-EPI) that covered the whole brain (in-plane resolution = 3.4 mm; 45 slices; slice thickness = 3 mm; slice spacing = 0.4 mm; TR = 2 s; TEs = 13, 32.3, 51.6 ms; flip angle = 70°; field of view = 220 mm; phase encoding = anterior–posterior; phase acceleration = 2; multiband factor = 3). Each run included 205 volumes (after excluding 5 dummy volumes), resulting in approximately 41 minutes of functional data per session and a total of ∼2.8 hours per participant across all sessions.

During each session, participants completed two runs of each of three passive viewing conditions. Parents and children within a family viewed the videos in the same fixed order within a session, but this order was randomized across families. All video content was novel across sessions, i.e., participants never viewed the same clips twice:

#### Narrative Movie

Scenes from the live-action movie *Dora and the Lost City of Gold* (Bobin, 2019) were shown in sequence. This film was chosen based on its appropriateness for a young audience, recent production (within the last decade), high engagement (Cutting et al., 2011), and a diverse cast.

#### Non-Narrative Clips

A compilation of short (24-65 seconds, mean = 46.4 sec) and highly viewed (> 1 million views) videos were sourced from *YouTube* and *TikTok.* These clips were chosen to maintain visual interest without providing significant narrative content. Footage included content such as Minecraft gameplay, stop-motion animation, TikTok dances, Rube Goldberg machines, and nature-based building videos. Background music was either retained from the original clip or added as instrumental tracks, with either minimal or no speech during the clips.

#### Low-Demand Video

Initially we planned to show Inscapes (Vanderwal et al., 2015) as a rest-like condition. However, during the pilot phase of this study, participants reported drowsiness during repeated viewings of the single Inscapes movie. Therefore, a custom condition was sourced from YouTube that featured calming and visually varied videos to better support the long and repeated scans. These videos included slow-moving scenes with instrumental music such as underwater wildlife, canyon walks, and aerial footage of Swiss and Japanese landscapes.

### Behavioral data collection

Post MRI, each participant completed surveys that assessed recall of the videos viewed during the six functional runs, along with self-reported measures of drowsiness and attention. Although these data were not analyzed as part of this study, participants reported remaining awake throughout scans, with a full analysis of these measures reported elsewhere (Rai et al., 2025).

### fMRI preprocessing

Preprocessing was implemented using a custom Nipype pipeline (v1.5.0) (Gorgolewski et al., 2011) with tools from FSL (v6.0.0) (Smith et al., 2004), ANTs (v2.3.4) (Avants et al., 2011), and AFNI (v21.1.16) (Cox, 1996).

For anatomical processing, T1-weighted images were processed using ANTs to perform bias field correction and skull stripping. Images were non-linearly registered to either the adult MNI 152 atlas or the pediatric NIHPD 7–11-year-old atlas (Fonov et al., 2009, 2011) based on each participant’s age. Tissue segmentations of gray matter, white matter, and cerebrospinal fluid were generated and warped back to native space. Following this, FSL was used to create tissue masks and AFNI was used to erode these masks by 7 levels.

For functional processing, each echo was processed separately. The first 5 volumes were discarded using FSL’s *ExtractROI* and motion correction was performed with FSL *MCFLIRT*, using the first volume of each run as the reference. Framewise displacement (FD) (Power et al., 2014) was computed from the second echo and used as the motion reference for subsequent steps. Slice timing correction was applied using FSL *slicetimer*, and co-registration across eachos was performed using FSL FLIRT, aligning echoes 2 and 3 to echo 1 for each run. Echoes were then combined using FSL *Merge*. All video runs within each session were then co-registered to the reference image (session 1, echo 1) using FSL *FLIRT*, and the six runs across conditions were concatenated using FSL *MERGE*.

### MEPI denoising and optimal combination

Using the *tedana* workflow (v0.0.12) (DuPre et al., 2021), pre-processed echoes were denoised and optimally combined via the T2* combination method (Posse et al., 1999). Next, PCA dimensional reduction was performed using the Akaike Information Criterion (AIC) option to improve ICA convergence. Independent component analysis (ICA) was used to classify based on the Kundu decision tree (v2.5) (Kundu et al., 2013) into BOLD (TE-dependent), non-BOLD (TE-independent), and uncertain (low-variance) components. Those automatically labelled as uncertain were then manually reviewed and re-classified as necessary to improve denoising accuracy, as suggested in prior work (Dipasquale et al., 2017; Griffanti et al., 2017; Mejia et al., 2020).

### Optimally combined MEPI processing

Subsequent processing was applied on the individual optimally combined (OC) data (4 sessions x 3 conditions x 2 runs). Each run was skull stripped using FSL *BET* and then FSL *FLIRT* for boundary-based registration to align to each participant’s T1-weighted image. We removed linear, quadratic, and mean trends, along with bandpass temporal filtering (0.01-0.08 Hz). We filtered high-frequency motion (>0.1 Hz) in the phase-encoding direction (Gratton, Dworetsky, et al., 2020), prior to nuisance regression (24 head motion parameters, white matter, cerebrospinal fluid, and the global signal). Volumes were then censored above a framewise displacement threshold of 0.15 mm (Power et al., 2012) as visual inspection suggested that there was an appropriate motion floor in these data. Lastly, runs were warped to native T1 space and concatenated together by viewing condition.

### Cortical surface projection

Cortical surfaces were generated from T1-weighted images using Freesurfer’s *recon-all* pipeline (v6.0) (Dale et al., 1999) and converted to Connectivity Informatics Technology Initiative (CIFTI) format (Glasser et al., 2013) via Ciftify’s *ciftify_recon_all* pipeline (Dickie et al., 2019).

### CIFTI fMRI data

Surface-based fMRI time series were generated via Ciftify’s *ciftify_subject_fmri* pipeline, using the ribbon-constrained sampling approach as in (Glasser et al., 2013) and smoothed with σ = 4mm geodesic kernel.

### Network identification in individuals via template-matching

A template-matching (TM) approach (Dworetsky et al., 2021; Gordon, Laumann, Adeyemo, et al., 2017) was used to generate individualized functional networks for each participant shown in Supplemental Figure 1 and 2. We opted for this approach over data-driven techniques (i.e., clustering algorithms such as Infomap or ICA) given the focus here was on comparing networks across age groups, and TM ensures this comparison is more reliable and direct (Dworetsky et al., 2021) while still reducing susceptibility to motion artifacts (Sylvester et al., 2023).

This approach requires a set of predefined network templates for individual-level matching. To avoid age-related bias in template creation, we created a combined child and adult network template using independent data. Specifically, group-level templates for adults (n = 384, mean age = 28.4 years; data = 60 min of resting-state fMRI; Dworetsky-HCP) and children (n = 38, mean age = 9.16 years; data = 26 min of resting-state fMRI over four runs; HCP-D) were obtained from the public MIDB Precision Brain Atlas repository (Dworetsky et al., 2021; Somerville et al., 2018). Adult and child network templates were derived from un-thresholded probabilistic maps (v1.0) based on the WashU 14-network atlas (Power et al., 2011). We binarized each child and adult network template by thresholding each network’s probabilistic map at 0.2. The final group-level maps for each network were then computed with a logical AND between child and adult templates. The resulting 14 network group-level consensus maps were labelled following the nomenclature from Power et al., (2011) as: default mode (DMN), frontoparietal (FP), visual (VIS), dorsal attention (DAN), ventral attention (VAN), salience (SAL), auditory (AUD), cinguloopercular (CON), somatomotor dorsal (SMd), somatomotor lateral (SMl), temporal pole (Tpole), medial temporal lobe (MTL), parietal memory (PMN), and parietal occipital (PON). The 14 network group-level templates were visualized using Connectome Workbench’s *wb_view* function (Glasser et al., 2013) and are shown in Figure 2.

To match each PreciseKIDS child and adult participant to this network template, vertex-wise dense FC matrices for the cortical surface were computed for each participant from 60 minutes of post-censored data (20 minutes of each viewing condition) using the Connectome Workbench function –*cifti-correlation.* Then, for each vertex, the FC profile was thresholded at the 95^th^ percentile, binarized, and compared against each network template by using the Dice similarity (Sørensen-Dice coefficient), defined as (Dice, 1945):

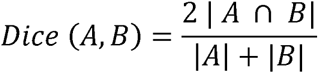

where A represents a vertex’s FC profile and B represents the group template.

Each vertex was assigned to the network with the highest Dice similarity, and continuous dice values were retained for analysis as described below.

Final individual-level networks were determined with a winner-take-all approach, where each vertex was assigned to the network with the highest Dice similarity; however continuous dice values were retained for further analysis as described below. Individual PreciseKIDS child and adult network maps were also combined to produce group-level network maps, by determining the most common network label within the child and adult groups (Figure 2). To facilitate comparison with other studies, we provide alternate network labels based on template matched Dice similarity computed using the Network Correspondence Toolbox (Kong et al., 2025) across four reference atlases: (Glasser et al., 2016; Gordon et al., 2016; Shirer et al., 2012; Thomas Yeo et al., 2011) as shown in Supplemental Table 1.

### Age group differences in network topography

Next, for each network we determined the overlap between individual-level adult and child network maps and thresholded by the number of participants at 0.2 for direct comparison to the HCP templates. We computed heat maps to indicate the number of participants in each group who were assigned a network label at a given vertex. Differences between groups were then visualized on the surface. Analyses in this study focused on 11 cortical networks: DMN, FP, DAN, VAN, SAL, CON, AUD, SMd, SMl, VIS, PON. The following 3 networks were excluded from analyses: Tpole, MTL, PMN, where appropriate. Tpole and MTL were excluded due to known signal dropout in anterior and medial temporal regions (Rai et al., 2024, 2025; Rua et al., 2018, p. 201), and PMN was excluded as it is more consistently detected with high-resolution imaging (Kwon et al., 2025). Group differences in network assignment at each vertex were assessed using mixed-effects logistic regression across networks, with age group as the main effect, controlling for fixed effects of sex and motion and random family ID intercepts. The resulting age group-effect beta coefficients were visualized with uncorrected p-values < 0.05 on the surface.

### Network assignment confidence differences

To assess whether there were group differences in the network assignment confidence, for each individual at each vertex, we computed the Shannon entropy (Shannon, 1948) of the normalized Dice similarity distribution defined as ():

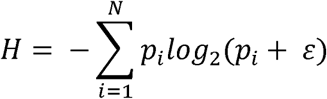

where p_i_ is the normalized Dice similarity for network *i* at a given vertex, and *N* is the total number of networks.

Higher entropy values indicate lower confidence, and lower values indicate greater confidence for a particular network assignment. Compared to the silhouette measure (Rousseeuw, 1987) which compares a vertex’s similarity to other vertices across communities (Tooley, Bassett, et al., 2022; Yeo et al., 2011), entropy provides a more computationally efficient estimate of how confidently a vertex belongs to its assigned network based solely on its own Dice similarity distribution.

We visualized entropy (as an index of confidence for network assignment) across the cortical surface. First, we tested whether children differed significantly from adults in subject-wise overall entropy (averaged across all vertices) using mixed effects models with fixed effects of sex and motion, and random family ID intercepts. We next applied this approach to vertex-wise entropy, where at each vertex, a mixed-effect model was used to examine the main effect of age group, while controlling for fixed effects of sex and motion, and random family ID intercepts. Standardized vertex-wise beta coefficients and uncorrected p-values < 0.05 were visualized on the cortical surface.

### Network surface area differences

We compared the cortical surface area occupied by each network in individualized child and adult network maps. Surface area of each vertex was computed separately for the left and right hemisphere using Connectome Workbench’s –*surface-vertex-areas* function. Then, for each network, we summed surface area values across network-assigned vertices and calculated proportions by dividing each network’s surface area by the total cortical surface area. We visualized the distribution of these percentages across networks and tested group differences in surface area for each network using a linear mixed-effects models with the main effect of age group while controlling for fixed effects of sex and head motion and random family ID intercepts.

Group differences were examined across all vertices and in a sensitivity analysis, only included thresholded vertices (Dice ≥ 0.3). This cutoff was chosen to exclude ambiguous network assignments, while retaining broad cortical coverage, and has been used in previous work (Boukhdhir et al., 2021). Across-network multiple comparisons correction was applied using the false discovery rate (FDR).

### Individualization of network topography

It has been suggested that development is associated with increasing individualization of the functional connectome (Demeter et al., 2025; Kaufmann et al., 2017; Li et al., 2025). Here, we asked whether network topography is more unique in low-motion children relative to low-motion adults (i.e., whether pairs of children are less similar than pairs of adults). Individualized network maps were used to compute Dice similarity between all pairs of children (excluding siblings) and all pairs of adults, for each network. The ciftiTools package was used to assess similarity on the CIFTI-format data (Pham et al., 2022) and R’s *lmer* function was used for linear modelling. To test for group differences in within-group similarity, we used a linear mixed-effects model with fixed effects for group (low-motion child-child vs. low-motion adult-adult), network, their interaction, and whether the pair shared the same sex. Random intercepts were added for each participant to account for the repeated measures across pairs as:

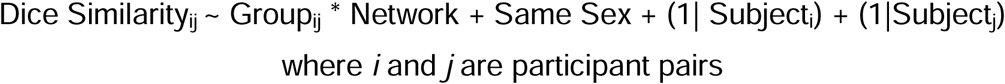

*Post-hoc* comparisons were performed to determine differences in within-group similarity per network with FDR-correction applied. Given prior evidence that adults achieved higher test-retest reliability than children when using high-probability regions of interest (Rai et al., 2025), we repeated analyses and used a linear mixed-effects model as described above with 30 minutes of adult data to approximate the reliability obtained by low-motion children with 60 minutes of data.

### Familial similarity in topography

Given prior evidence on the heritability of the connectome (Miranda-Dominguez et al., 2018), we examined whether relatedness resulted in greater similarity in network topography. We compared related child-adult pairs (i.e., child-parent) against unrelated child-adult pairs. First, we computed vertex-wise Dice similarity between all possible child and adult individualized network map pairs (n = 576 pairs across 24 children x 24 adults) for each of the 11 cortical networks. Group differences in Dice similarity were tested with linear mixed-effects models, including main effects of group relation (two-level variable of related vs. unrelated) and network, fixed effects of same sex pairing, and random intercepts for each participant to account for repeated pairings as:

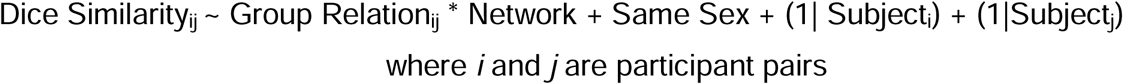

We applied the same linear mixed-effects model to compare Dice similarity for each network separately between related and unrelated pairs, defined as:

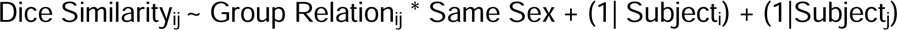

### Age effects on dense within-and between network FC: impact of individualized networks

To complement the above network topography analyses, we examined age group differences in within- and between-network FC, using both individualized networks and the PreciseKIDS group templates. Dense FC matrices were computed using the same 60 minutes of post-censored data used for template-matched network creation. In a separate validation analysis, an additional 15 minutes of post-censored data (5 minutes of each condition) were analyzed to assess results when data used to calculate functional connectivity are independent of the data used to define networks. We applied vertex-wise distance censoring to each vertex’s FC value using the Conte-69 surface mesh (Van Essen et al., 2012). All FC edges within 30 mm were excluded as in (Demeter et al., 2023) to reduce spatial autocorrelation (Jeganathan et al., 2024). FC values were then Fisher z-transformed and averaged across vertices using three network approaches: 1) group-level winner-take-all templates applied to individuals 2) Individualized network maps derived from template-matching, and 3) high-confidence individualized network maps that included only those vertices with a Dice similarity ≥ 0.3.

We compared age groups on each within- and between-network FC measure using linear mixed-effects models with the main effect of age group, controlling for fixed effects of sex and head motion, and random family ID intercepts. Multiple comparisons correction was applied using the false discovery rate (FDR) and effect sizes are shown as the standardized beta coefficients from the model. We noted differences in significant findings across network approaches.

## Results

### Age group differences in network topography

Templates derived from adult HCP and child HCP-D data provide a point of reference for potential expected age differences in topography. Comparing the thresholded (i.e., network was present in 20% of the sample) adult HCP and child HCP-D network maps (Figure 1A), we observed that all large-scale cortical networks had regions of convergence (overlap) and regions of divergence (no overlap) between age groups. Specifically, DMN, SMd, SMl, and VIS networks showed greater cortical coverage in adults compared to children, while CON, DAN, VAN, and PON networks showed greater coverage in children.

**Figure 1:**
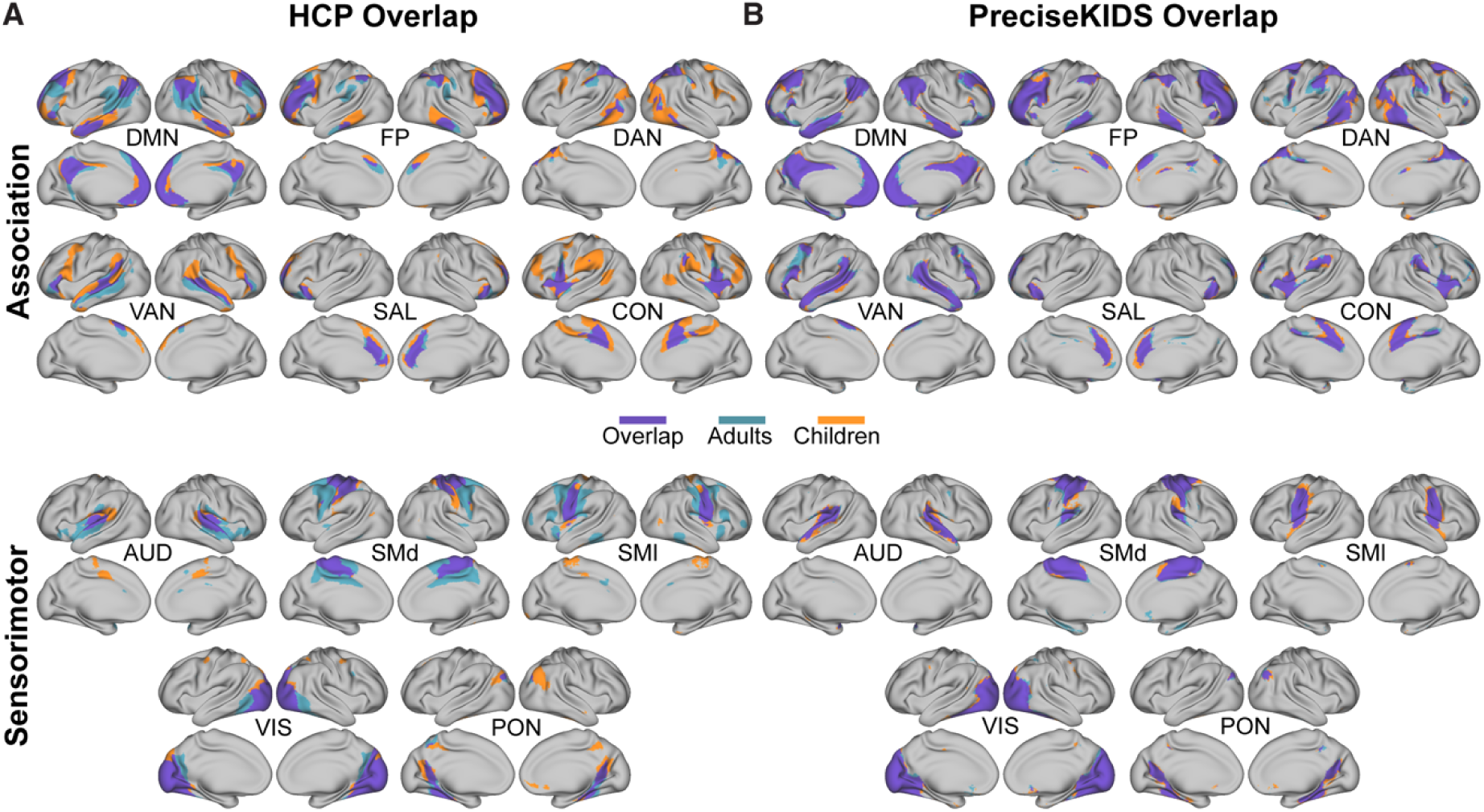
Adult and child cortical networks in HCP/HCP-D and PreciseKIDS. A) Spatial overlap maps derived from the HCP/HCP-D datasets, thresholded at ≥ 0.2 show both overlapping networks and regions of divergence across age groups. Adults exhibit expanded coverage in DMN, SMd, SMl, and VIS networks, while children show broader coverage in CON, DAN, VAN, and PON. B) Overlap maps derived from the PreciseKIDS dataset thresholded at ≥ 0.2 are broadly similar network architecture to HCP/HCP-D derived templates and show high convergence between children and adults.

Next, we considered networks identified in our data and applied the same threshold (0.2) to the individualized networks (Figure 1B). In our data, relative to the HCP/HCP-D data, we found greater convergence (overlap) of network maps between children and adults, with subtle differences in network boundaries. The remaining networks (Tpole, MTL, and PMN) are shown in Supplemental Figure 3. When derived from the winner-take-all maps, the PreciseKIDS networks encompassed more of the cortical surface and showed a reduced PMN, and expanded FP, DAN, and VAN, relative to the HCP winner-take-all maps in Figure 2. Of note, relative to the HCP/HCP-D atlases, which are derived from resting-state data, PreciseKIDS networks show expanded overlap in DAN, VAN, and AUD regions, which may reflect the naturalistic stimuli. We also note that network labelling can vary across commonly used atlases (e.g., Glasser et al., 2016; Gordon et al., 2016; Shirer et al., 2012; Thomas Yeo et al., 2011) and provide alternative network labels to facilitate comparison with prior studies in Supplemental Table 1. Taken together these observations suggest that inferring topography age effects using data collected with different parameters should be done with caution, and that for the age range considered here, age effects on topography are subtle.

**Figure 2:**
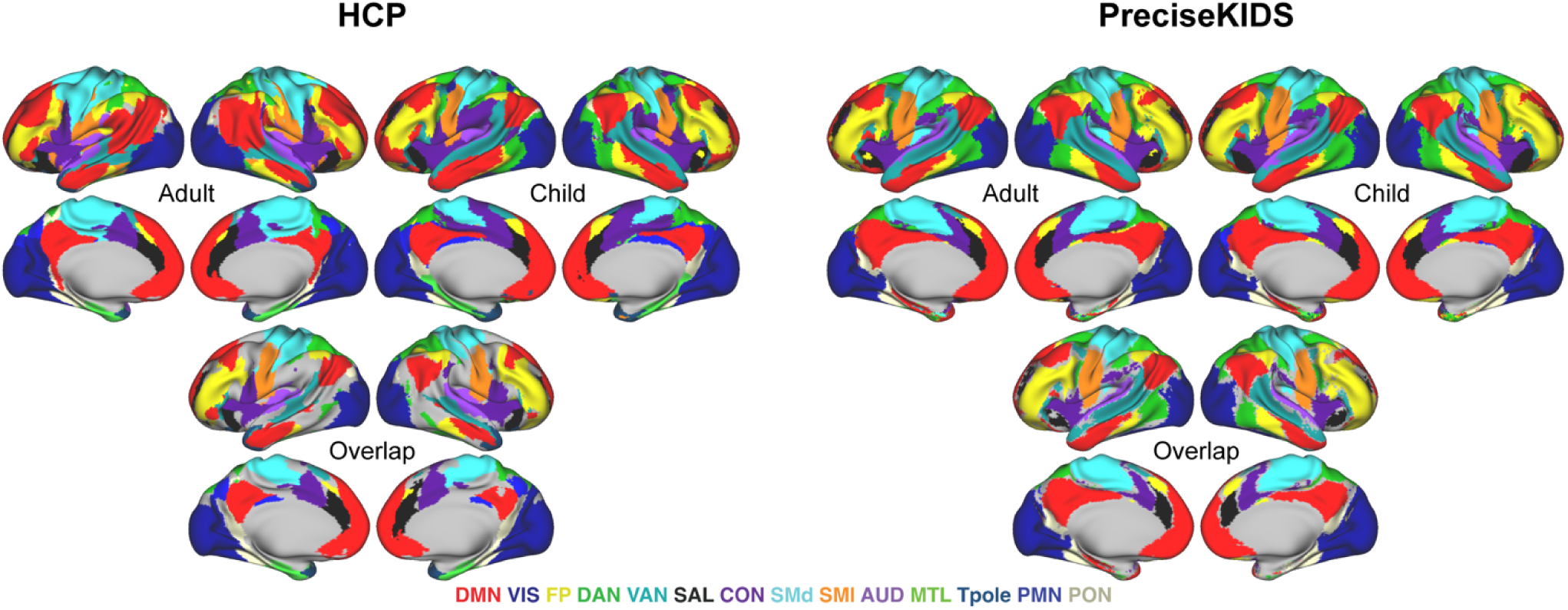
HCP and PreciseKIDS winner-take-all networks. In the left panel, thresholded winner-take-all adult (HCP), child (HCP-D), and the overlap between them is shown using the HCP datasets. In the right panel, PreciseKIDS winner-take-all adult, child, and overlap surface maps are shown. Compared to HCP, the PreciseKIDS networks showed a reduced PMN and an expanded FP, DAN, and VAN.

Vertex-wise network assignment density for children and adults is shown in Figure 3. For most networks there are ‘core’ regions where all 24 individuals in each age group had overlapping assignment (red in Figure 3), whereas network boundaries displayed more heterogeneous assignment patterns across individuals, with some vertices assigned to a network in only a small number of individuals (blue regions in Figure 3). Regions that showed idiosyncratic assignment patterns (i.e. vertices where a small proportion of the sample was assigned to a given network) occurred at similar rates across both children and adults reflecting individual differences in network organization rather than age-related effects (Figure 3). For example, a similar proportion of the child and adult samples showed a temporo-parietal region that associated with the PON and regions of the posterior cingulate cortex and precuneus that associated with the FP network. Calculating surface area of each network density map – a measure sensitive to the spread of a network across individuals – showed that child network assignments covered a larger surface area for nearly all networks compared to adults, with the most pronounced differences for the DAN, VIS, FP, VAN, SMd, SML, and CON networks (Supplemental Table 2). The SAL network was the single network that exhibited a larger spread in adults compared to children. This suggests that children’s network territories are more variable across individuals and/or more spatially diffuse.

**Figure 3:**
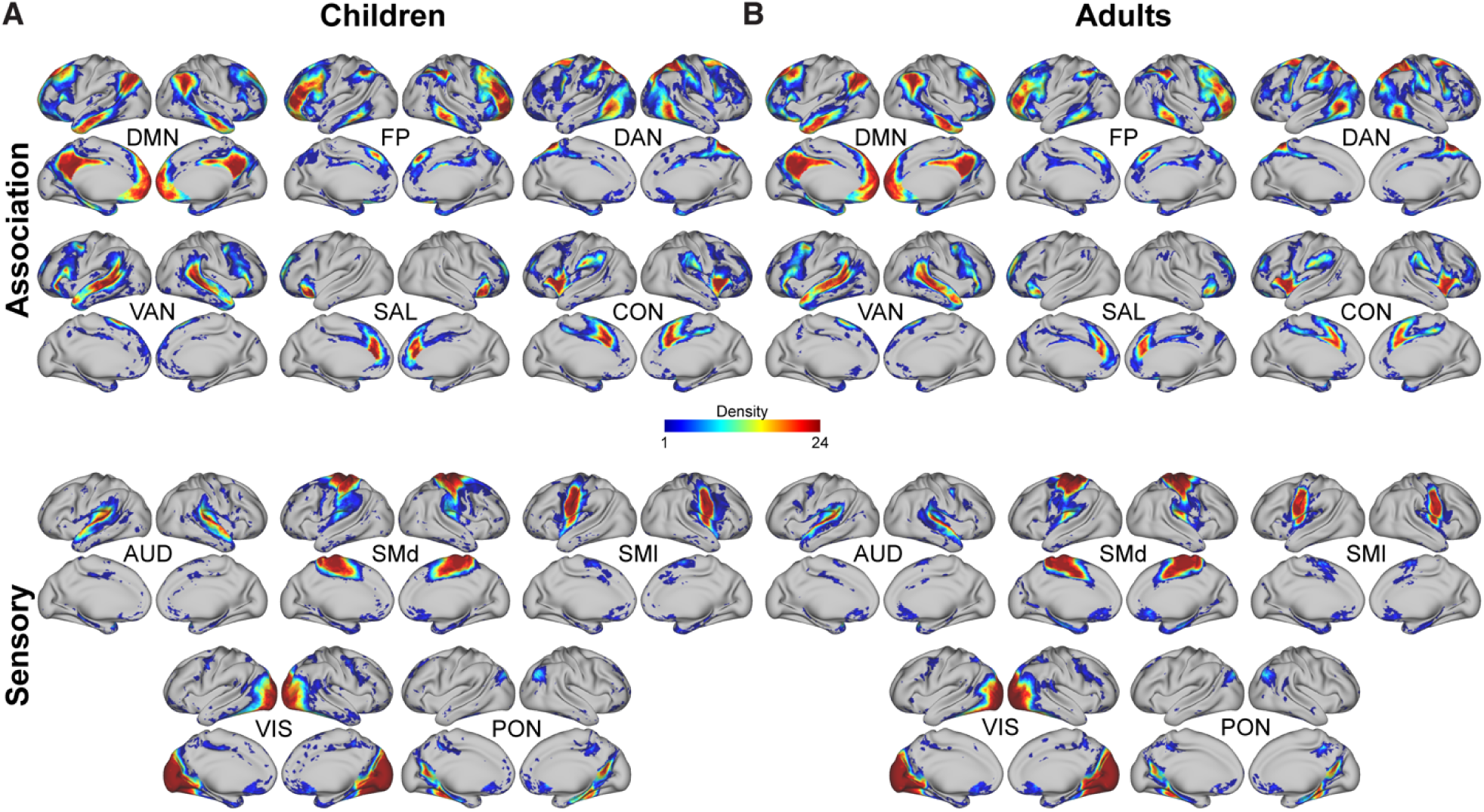
Vertex-wise network assignment density between children and adults. Group density surface maps for each network across (A) children and (B) adults.

We asked whether there were consistent shifts in network borders between children and adults, by conducting between-group vertex-wise comparisons of network membership (Figure 4A). The networks with the largest contiguous sets of significant vertices (i.e., age group effect exceeding p < 0.05 uncorrected) included SMd (largest cluster size = 208.05 mm^2^), PON (largest cluster size = 152.48 mm^2^), CON (largest cluster size = 149.16 mm^2^), and SAL (largest cluster size = 100.37 mm^2^). These clusters were largely found at network borders, for example at the dorsal somatomotor network boundary; however certain clusters existed within core regions for CON and SAL networks (Figure 4B). The networks with the greatest effect of motion included DMN, FP, SMd, and PON (p-uncorrected < 0.05) and the networks with largest sex effects included FP, DAN, VAN, and SAL (p-uncorrected < 0.05) (Supplemental Figure 4).

**Figure 4:**
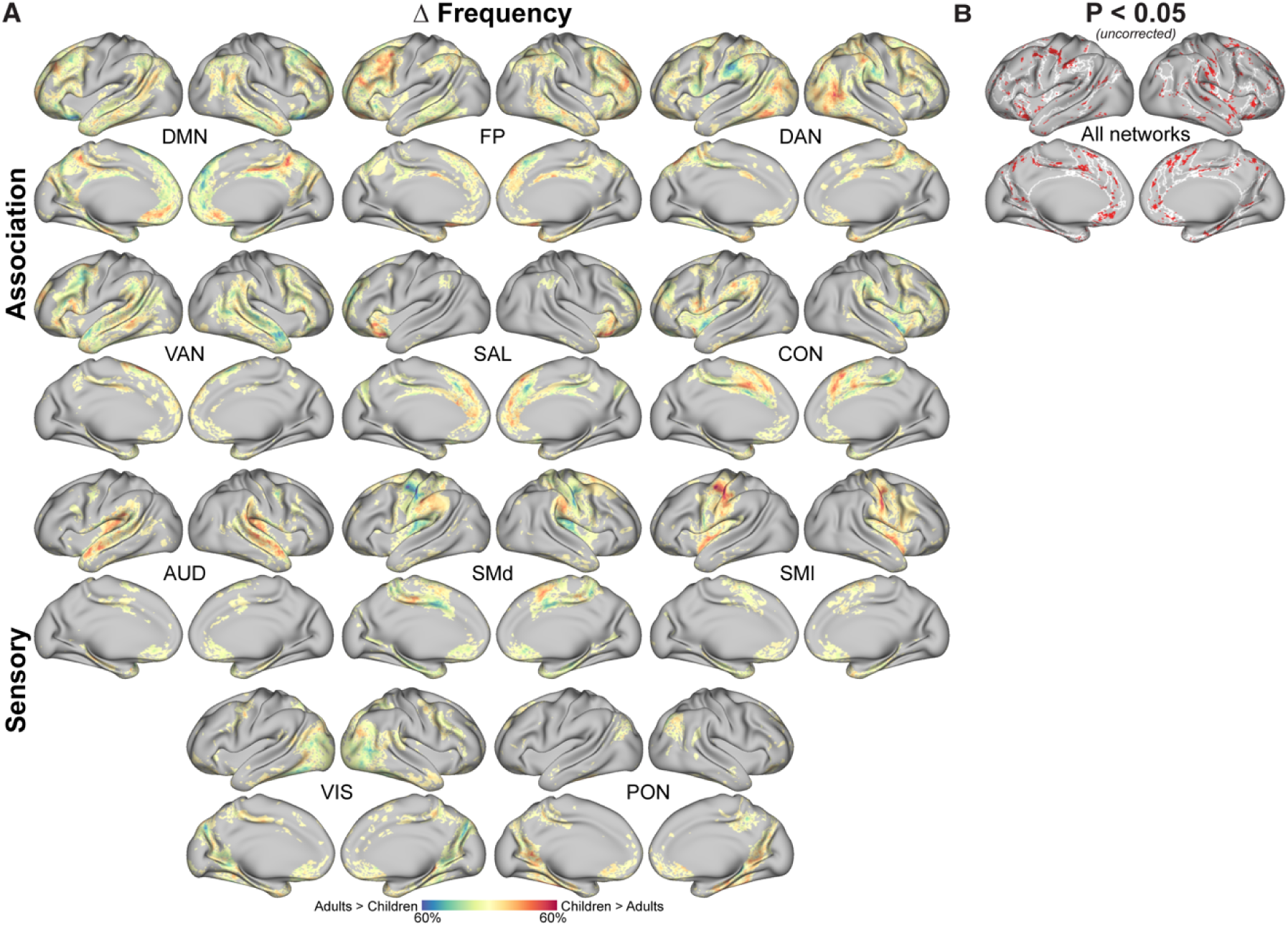
Vertex-wise differences in network assignment frequency between children and adults. A) Proportional group differences in network assignment are shown, with warm colors indicating more frequent assignment in children, and cool colors indicating more frequent assignment in adults. Maximum values indicate a given network assignment was observed in 60% of participants within groups. B) Vertices with between-group differences (p < 0.05 uncorrected, mixed logistic regression) are shown in red, with the winner-take-all network assignments for SAL, CON, SMd, and PON outlined in white.

### Network assignment confidence differences

To evaluate age effects on network assignment confidence, we computed individual vertex-wise Dice coefficient entropy, which were then averaged within groups for each vertex (Figure 5). Network assignment confidence was highest in regions of the visual and motor cortices (Figure 5A) and lowest in high dropout regions (temporal pole and orbitofrontal cortex). In association regions there was relatively higher confidence in core regions of the DMN and relatively lower confidence in boundaries of association networks, such as the right temporoparietal junction (Figure 5A). Averaged entropy values across all vertices for each individual showed that children (mean = 2.200, s.d. = 0.131) had significantly higher overall entropy relative to adults (mean = 2.012, s.d. = 0.093) (β = 0.108, *z* = 3.604, p < 0.001) and that entropy significantly associated with motion (β = 0.000, *z* = 12.026, p < 0.001), but not sex (p = 0.566). Vertex-wise entropy comparisons showed many regions where entropy was higher in children (lower confidence) relative to adults (positive β; Figure 5B). When vertex-wise entropy maps were thresholded at p < 0.05 uncorrected (Figure 5B), children showed lower confidence in larger clusters within the boundaries of DMN (largest cluster size = 329.06 mm^2^), SMd (largest cluster size = 576.54 mm^2^), and AUD (largest cluster size = 468.47 mm^2^). Adults showed lower confidence only in one region, along the border of the VIS network (largest cluster size = 695.09 mm^2^).

**Figure 5:**
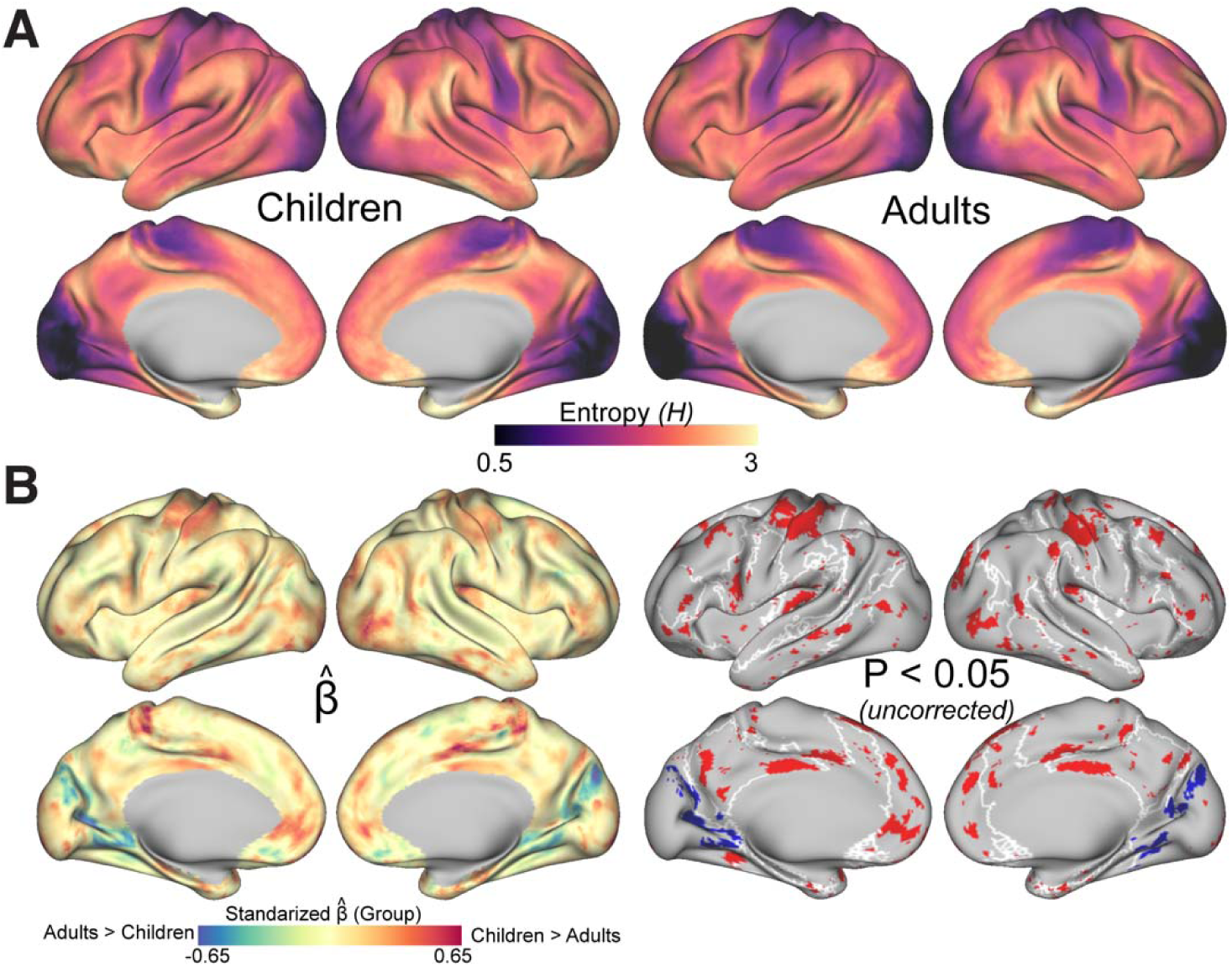
Network assignment confidence differs between children and adults. A) Vertex-wise surface maps show the entropy of dice coefficients across networks for children and adults. Lower entropy values (dark purple regions) indicate higher confidence in network assignment, whereas higher entropy (yellow regions) reflect less certainty. B) The left panel shows the standardized regression coefficients from the mixed-effects model contrasting between children and adults. Warm colors reflect greater entropy in children relative to adults (less confidence in children) and cooler colors indicate greater entropy in adults (less confidence in adults). Right panel shows vertices with uncorrected p < 0.05 group differences in red for children and blue for adults. DMN, SMd, AUD, and VIS networks are outlined in white.

### Network surface area differences

For each network, we compared surface area between children and adults and found that, PON and SMl were relatively smaller in adults (p-uncorrected < 0.05), however this did not survive FDR correction (pFDR > 0.05) (Figure 6; Supplemental Table 3). Head motion was associated with surface area for the DAN and SAL networks (p-uncorrected < 0.05), with only the SAL network significantly impacted by motion after correction (pFDR = 0.005). No significant effects of sex were observed across any networks after correction (pFDR > 0.05). Findings were comparable when analyses were restricted to high-confidence regions (Dice ≥ 0.3; Supplemental Figure 5; Supplemental Table 4).

**Figure 6:**
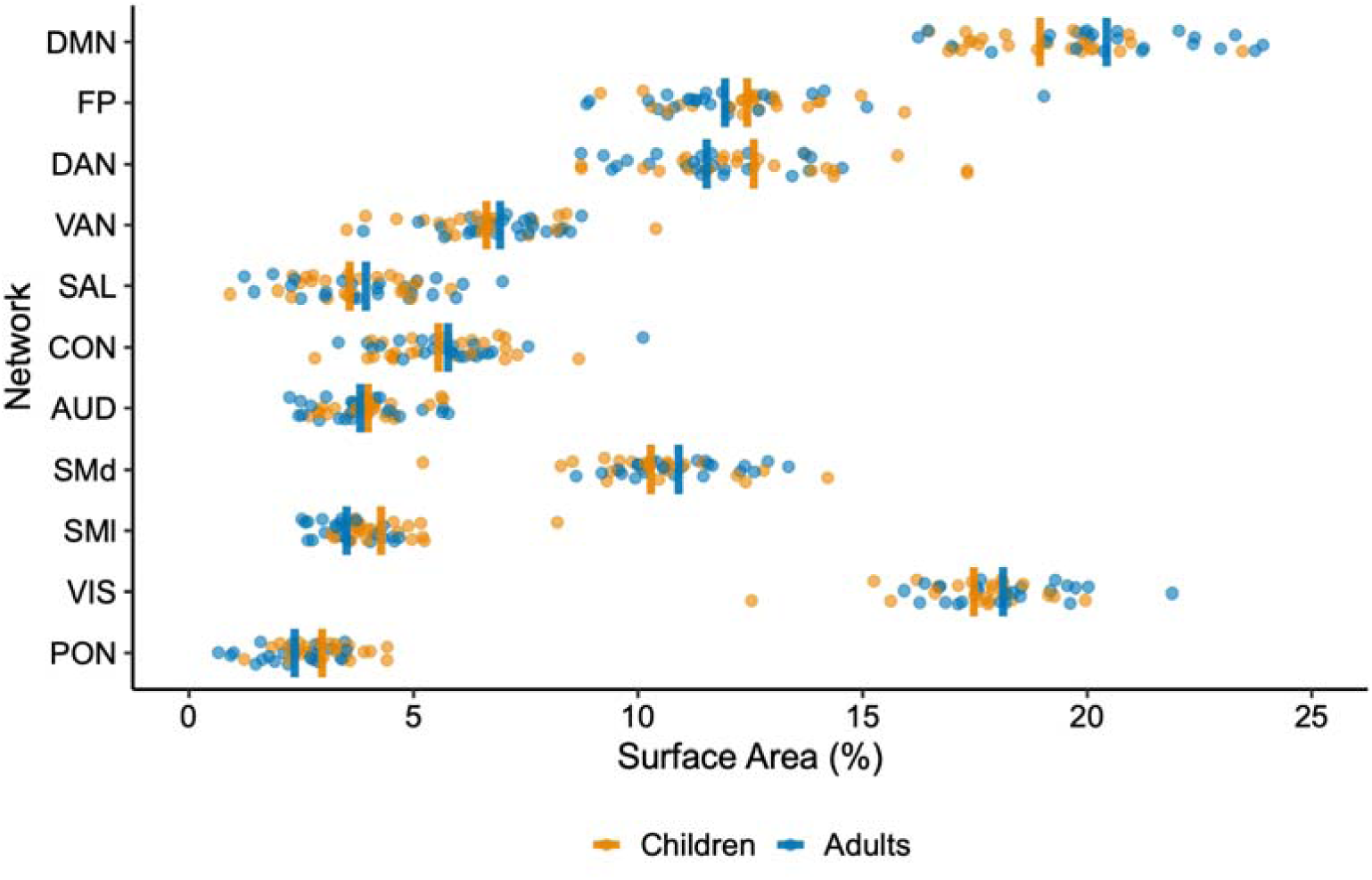
Surface area comparisons of functional networks between children and adults. Surface area (% of total cortical surface area in each individual) values are shown across networks in children (orange) and adults (blue), with group means. We found no significant differences between groups, controlling for sex and head motion.

### Individualization of network topography

We next asked whether there is increased individualization of network topography in low-motion adults (LMA; n = 22) relative to low-motion children (LMC; n = 10). To this end, we asked whether network topography is more individualized (i.e., less similar, on average) between pairs of adults relative to pairs of children (Figure 7). Within-group similarity in network topography showed no significant overall effect of age group (p = 0.107), a significant effect of network (all p < 0.002) and no effect of same/opposite sex pairs (p = 0.638). There were significant interactions between networks and age group, using DMN as the dummy-coded reference network. We followed up with *post-hoc* comparisons which showed that LMA pairs had greater within-group similarity (i.e., less within-group individualization) compared to LMC pairs for DAN (β = 0.049, *z* = 3.42, pFDR < 0.001), CON (β = 0.071, *z* = 4.985, pFDR < 0.001), and SMd (β = 0.083, *z* = 5.80, pFDR < 0.001). LMC pairs had significantly less individualization (greater similarity) than LMA pairs only for the SAL network (β = −0.105, *z* = −7.33, pFDR < 0.001) and differences were not significant for remaining networks (Supplemental Table 5). To ask whether these results are driven by greater reliability in adults (Rai et al., 2025) we repeated this analysis using 30 minutes of post-censored data in adults and 60 minutes in children. We found comparable results (Supplemental Figure 6), suggesting that greater reliability in adults is unlikely to be driving this effect.

**Figure 7:**
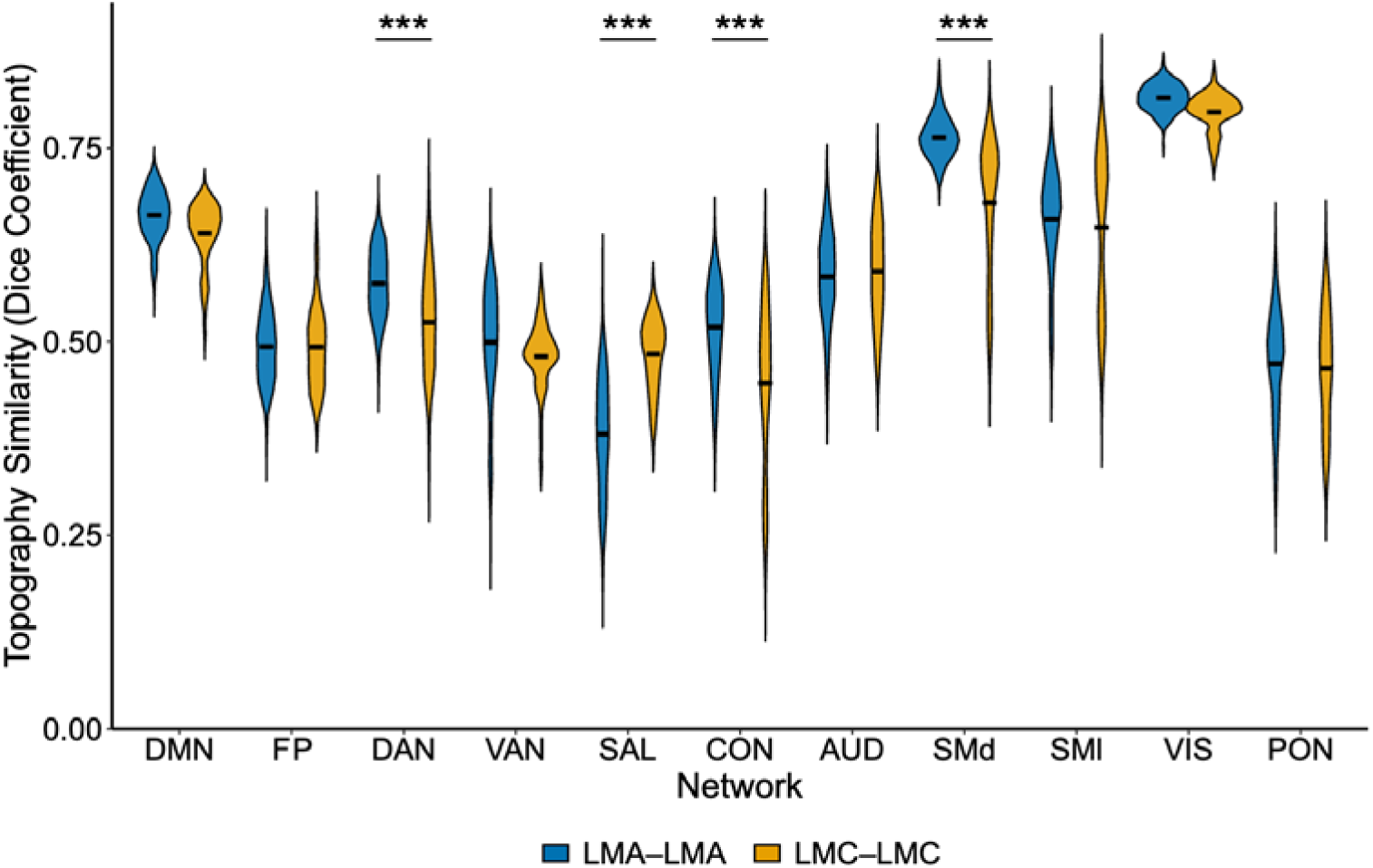
Within-group similarity of network topography among low-motion adults and low-motion children. Pairwise dice similarity in network assignment for low-motion adults (LMA-LMA; blue) and low-motion children (LMC-LMC; orange) for each network. There was a significant effect of network (p < 0.001) and interaction between network and group (p < 0.05). Asterisks denote *post-hoc* comparisons with FDR correction: *** p < 0.001.

### Familial similarity in topography

As this dataset included parent-child pairs, we asked to what extent network topography is similar within families. In an omnibus test across all networks, we found that the overall effect of relatedness was not significant (p = 0.33; Figure 8) across all networks. There was a significant effect of network (all p < 0.001), and no significant effect of sex (p = 0.605) or interaction between network and relatedness (all p > 0.05). On a network-wise level, however, we found significantly greater network topography similarity for related pairs over unrelated pairs across multiple networks: DMN (β = − 0.013, *z* = −2.88, pFDR = 0.015), FP (β = −0.021, *z* = −3.30, pFDR = 0.006), DAN (β = − 0.022, *z* = −3.62, pFDR = 0.004), and VAN (β = −0.017, *z* = −2.41, pFDR = 0.044).

**Figure 8:**
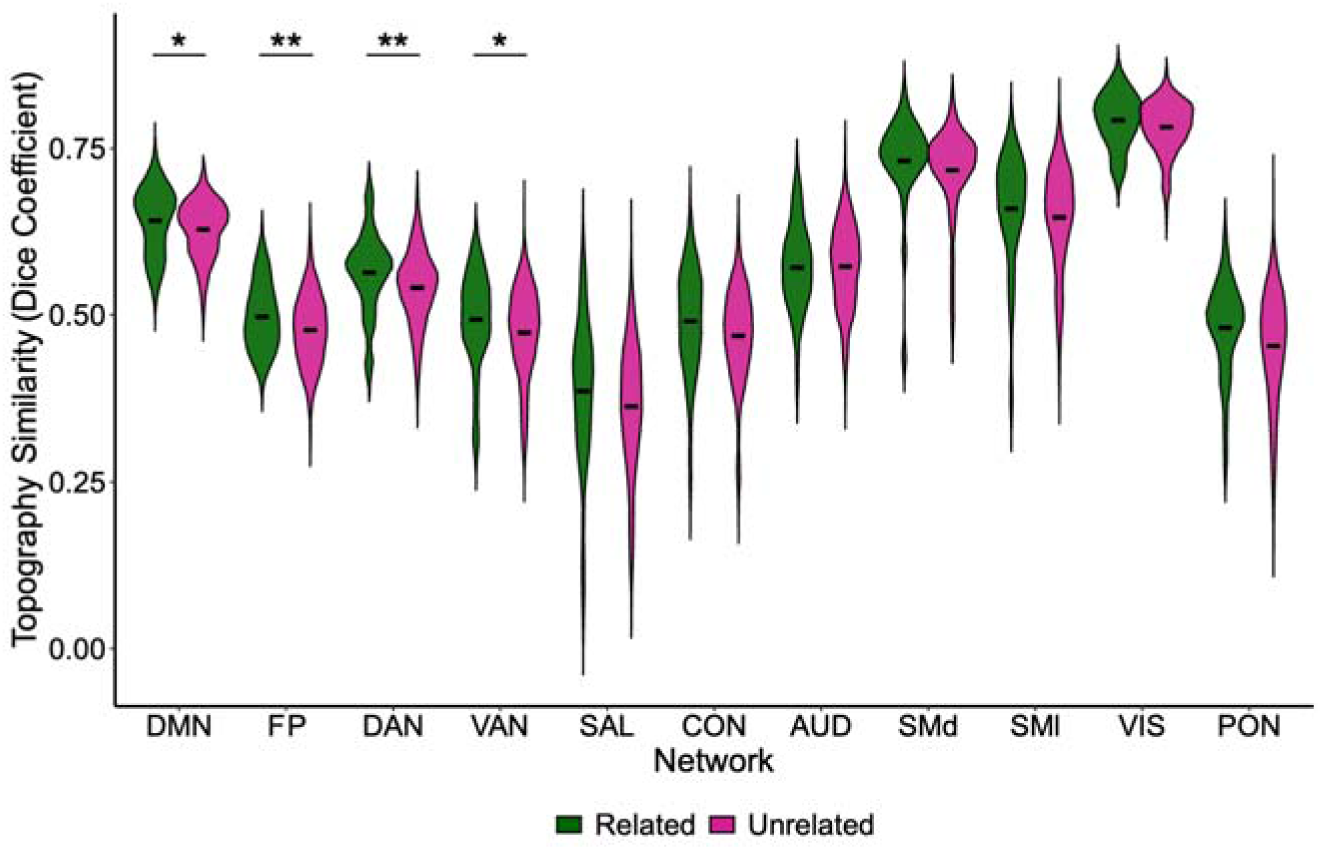
Network topography similarity in related versus unrelated pairs. Pairwise Dice similarity in network assignment is shown for related pairs (green) and unrelated pairs (magenta) across networks. Overall relatedness effects were non-significant, though network-wise comparisons showed greater similarity for related pairs across DMN, FP, DAN, and VAN networks. Significance asterisks denote network-wise comparisons with FDR correction: * p < 0.05, ** p < 0.01.

### Age effects on dense within-and between network FC: impact of individualized networks

Finally, we considered age effects on within- and between-network FC, and the impact of individually defined networks on group differences. We compared FC measures derived with the PreciseKIDS group network maps, individualized network maps, and individualized network maps thresholded to include only high-confidence vertices (Dice ≥ 0.3). Across all three approaches, adults showed stronger within-network FC across most networks compared to children (Figure 9A), and effects were most pronounced within and between SMd and SMl networks (β range = 0.051 to 0.143). Other strong adult>child within-network effects included DMN-DMN (β range = 0.029 to 0.067), DAN-DAN (β = 0.056 to 0.057), and VAN-VAN (β = 0.041 to 0.070). In a separate analysis using 15 minutes of independent data for calculating FC (i.e., distinct from the data used for template-matched networks), we observed similar results, with adults showing greater within-network FC relative to children (Supplemental Figure 7).

**Figure 9:**
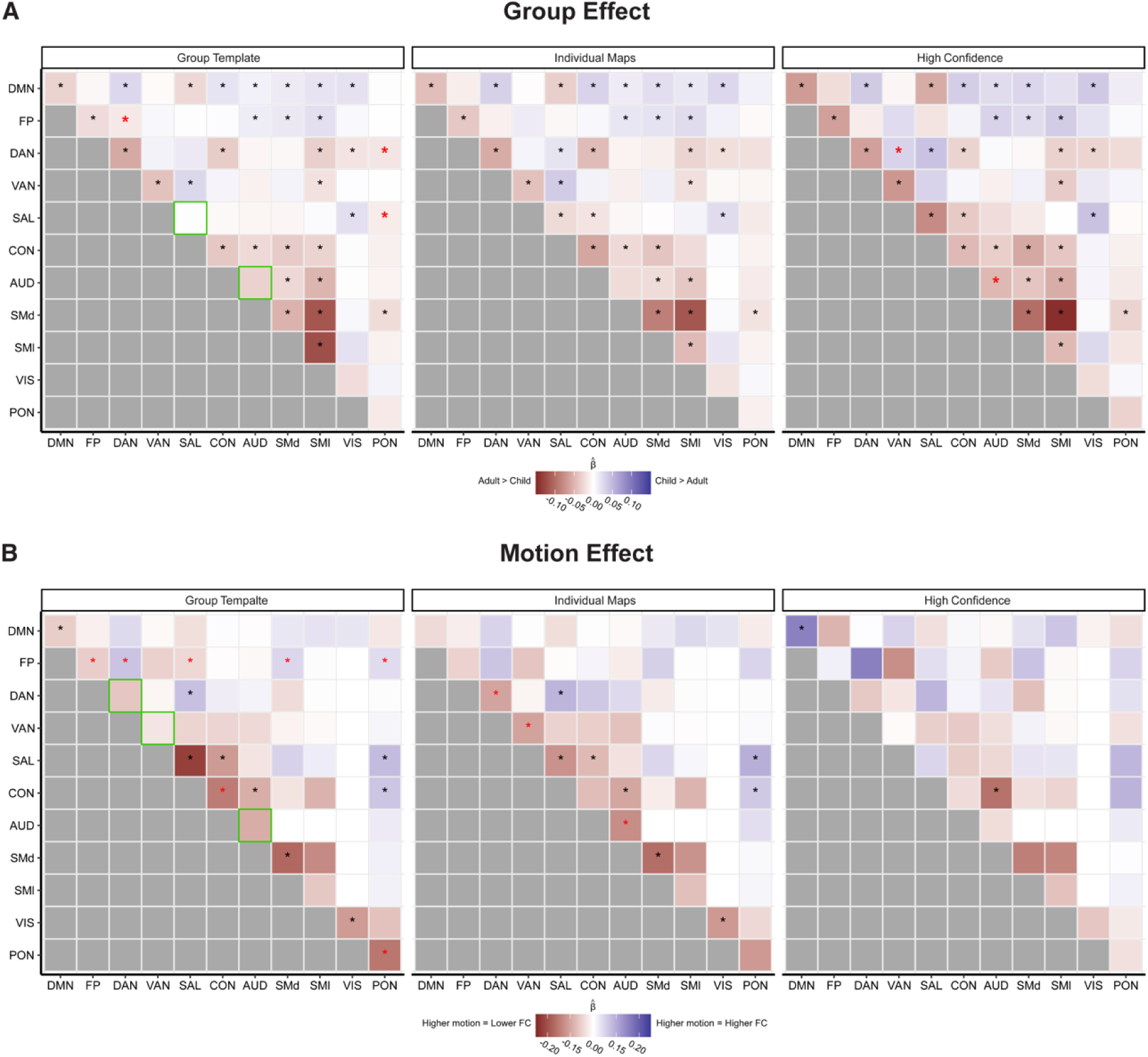
Differences in FC across networks and mapping approaches between children and adults. Standardized beta coefficients from linear mixed-effects models are shown for analyses using group-level templates (left), individualized networks (middle), and high-confidence individualized (Dice ≥ 0.3) (right) approaches. A) Beta coefficients from the group effect indicate stronger FC in adults relative to children in red and stronger FC in children relative to adults in blue. Across all approaches, adults exhibited stronger within-network FC than children. B) Shades of red indicate motion effects where higher motion decreased FC estimates (negative betas), whereas shades of blue indicate motion increased FC estimates (positive betas). Significance asterisks denote comparisons with FDR correction (p < 0.05). Red stars reflect significance for a specific approach only. Green boxed outlines indicate the difference in within-FC between the group template and either of the individualized approaches.

Beyond commonalities, individual and high-confidence mapping showed effects that are suggestive of both false positives and false negatives in the group template approach. Specifically, certain age group effects on between-network FC (e.g., FP-DAN, FP-PON, SAL-PON) were significant only when using the group network map, but these patterns were not significant when using the individual or high confidence network maps. Additionally, specific age group effects on within-network FC detected with individualized network maps (e.g., SAL-SAL for both individualized maps and AUD-AUD for the high-confidence maps) were not significant with the group network map. The individual mapping approach showed no uniquely significant effects, while the high confidence approach showed unique differences between DAN-VAN and AUD-AUD networks. There were significant effects of head motion in a majority of edges using the group template approach, with the largest effects for SAL-SAL, SMd-SMd, CON-CON, and PON-PON (Figure 9B). Effects were reduced using the individualized approach and only found within-networks for DAN-DAN, VAN-VAN, and CON-CON edges. The high-confidence approach further diminished motion effects, where significant motion effects were only detected for DMN-DMN (inflated FC due to motion) and CON-AUD (reduced FC due to motion) edges. These differences demonstrate the utility of individual mapping for ascertaining robust group differences in FC.

## Discussion

In this study, we collected a unique dataset featuring deeply sampled fMRI from parent-child pairs using passive viewing. We used these data to assess age effects on vertex-wise shifts in network assignment, assignment confidence, network surface area, factors impacting inter-individual topography similarity, and the impact of individualized network mapping for estimating age effects on FC. Our findings showed that while large-scale network architecture is broadly shared across age groups, children exhibit lower overall confidence in network assignments and weaker within-network FC relative to adults. We further found that age effects on FC differed when using individualized network mapping relative to a group network map, highlighting the potential for individual-specific network definitions to improve specificity in developmental studies.

Using individually defined functional networks, we found that large-scale network topography is broadly similar between pre-adolescent children and adults with only small age-group differences in assignment frequency, despite inter-individual variation in both age groups. While these findings are consistent with other work showing general similarity in network topography from relatively early in childhood through adolescence (Tooley, Park, et al., 2022; Tu, Myers, et al., 2025), we also found fewer group differences than suggested in other work (Sun et al., 2025; Tooley, Bassett, et al., 2022). Specifically, comparing ABCD and HCP samples, Tooley et al. (2022) found expanded limbic and visual territory and contracted default and frontoparietal network territory in children compared to adults. A mega-analysis pooling data across many samples also suggested some differences in network assignment of ventral attention and somatomotor regions when contrasting across age groups (Sun et al., 2025). Differences in findings from our work may reflect that pooling across datasets using different protocols can conflate age and dataset effects, underscoring the need for studies covering wide age ranges with harmonized protocols, as well as longitudinal work to capture within-individual change.

We did not observe significant differences between children and adults in cortical surface area percentages for any networks and age effects at network boundaries were small and would not survive vertex-level multiple comparisons. With those caveats, we note that age effects were most prominent along dorsal somatomotor and parieto-occipital network boundaries, and within core regions of the cingulo-opercular and salience networks. Differences observed in sensory and motor regions suggest that though these networks mature early relative to association regions, they may nonetheless undergo continued topography refinement (Fair et al., 2009; Koolwijk et al., 2024; Satterthwaite et al., 2013; Supekar et al., 2009) and cortico-subcortical connectivity maturation across development (Badke D’Andrea et al., 2023; Greene et al., 2014). We also note these effects may depend on network definitions and that alternative parcellations, such as the three-network sensorimotor model (Demeter et al., 2025; Gordon, Laumann, Gilmore, et al., 2017) or an inter-digitated action network (Gordon et al., 2023), could impact findings. Markedly, one of the largest differences in network assignment confidence between children and adults emerged for the somatomotor network, warranting careful interpretation of the vertex-wise network assignment differences in the SMd network regions. Regarding effects within regions of the cingulo-opercular and salience networks, this finding aligns with prior work suggesting that these networks exhibit continued strengthening and refinement with age (Marek et al., 2015). Importantly, although cross-sectional age effects can be suggestive of changes, longitudinal data are needed to assess developmental stability, or change, in individual topography (Lynch et al., 2023).

Network assignment entropy analyses showed that children exhibit lower confidence (i.e., higher entropy) in network assignments than adults, particularly at the boundaries of the default mode, dorsal somatomotor, auditory, and visual networks. This extends prior work by Tooley et al. (2022), who showed lower overall confidence in children’s functional organization compared to adults. Together our results suggest that functional networks do not undergo large boundary shifts between pre-adolescence and adulthood, but that connectivity refinement continues and contributes to increased confidence. The exception was reduced confidence at visual network borders in adults compared to children, an unexpected finding that may reflect age-related differences in response to passive viewing videos. Importantly, regions with lower confidence in adults have previously been interpreted as integration zones or ‘hub’ regions that support cross-network communication (Dworetsky et al., 2021). Hence under this framework, higher entropy or lower confidence in children found in this study could reflect increased functional flexibility rather than immature network organization or reduced functional specialization.

Comparisons of different methods for defining functional networks (group vs. individual-levels) in terms of impact on functional connectivity analyses demonstrated the importance of network definition approaches. While all three approaches to defining networks (group template, individualized, and high-confidence individualized) showed that adults have stronger within-network connectivity than children, the specific patterns of significant differences varied substantially across approaches. Age effects echo previous work showing increased network segregation and functional specialization with age (Fair et al., 2009; Supekar et al., 2009; Tooley, Park, et al., 2022). The group template approach also showed several between-network connectivity differences that were absent when using individualized approaches and had the greatest impact on connections from head motion. In contrast, the high-confidence approach isolated only three motion-sensitive functional connections (e.g., DMN-DMN, SMl-AUD, CON-AUD), underscoring the relative robustness of individualized approaches to motion. Given that the high confidence approach excludes lower confidence regions, these may represent false positives due to misalignment between group templates and individual network boundaries, especially in developmental studies (Tu, Wang, et al., 2025). Collectively, these results indicate that individualized methods offer improved specificity and sensitivity to variation in FC (Dworetsky et al., 2021; Hermosillo et al., 2024).

Heritability of FC and network topography has been demonstrated by prior twin studies (Anderson et al., 2021; Busch et al., 2024; Colclough et al., 2017; Demeter et al., 2020; Glahn et al., 2010; Miranda-Dominguez et al., 2018; Sinclair et al., 2015). Extending on these studies, we found greater similarity in DMN, FP, DAN, and VAN topography in parent-child pairs than in unrelated adult-child pairs. We note that our design may capture both shared genetics and environmental effects (Boomsma et al., 2002), which are more separable using heritability designs. Parent-child pairs also span different developmental stages, potentially introducing age-related variance that may mask overall genetic similarity. Our findings add to the current literature suggesting familial relatedness is a potential source of individual variation in network topography (Anderson et al., 2021) and underscore the need for larger, genetically informed precision samples to robustly characterize the heritability of the functional network architecture (Colclough et al., 2017; Ge et al., 2017).

Network assignment density maps – maps of the number of child or adult participants with a given network assignment at each vertex – showed greater spread for nearly all networks in children (except for the salience network). Network size was not larger in children, when assessed by surface area. Together this suggests that topography in children is more individually unique, a finding generally supported by pairwise similarity analyses, which showed a predominant pattern of higher pairwise similarity in adults. These findings are somewhat contradictory with prior literature suggesting that functional connectivity becomes more individualized with increasing age (Demeter et al., 2025; Kaufmann et al., 2017), although no prior studies to our knowledge have looked at individualization of topography, focusing instead on dense (Demeter et al., 2025) or parcellated (Kaufmann et al., 2017) connectome individualization. The measure used here, dice similarity of network topography, captures one aspect of individualization – dissimilarity from others – and not the other contributor to fingerprinting techniques which is self-similarity. Some prior work has suggested that increasing individualization with age is primarily driven by increased self-stability (Graff et al., 2022; Kaufmann et al., 2017), with a smaller impact of self-other distance. There are additional methodological considerations that may contribute to different findings across studies. First, we used naturalistic viewing instead of resting-state scans, which may have reduced individualization effects due to shared stimulus-driven activation patterns (Finn et al., 2017; Vanderwal et al., 2017, 2021). Additionally, although video content was similar, we used different clips across runs, which could have attenuated stable individualization effects. Reliability may also be a factor. Here, we used 60 minutes of fMRI per person, while some other work has used upwards of 90 minutes of low-motion data (Demeter et al., 2025; Lynch et al., 2023). Finally, although we used stringent motion censoring and denoising approaches, residual motion effects, especially in higher motion children, could still bias group differences observed (Rai et al., 2025).

Strengths of this study included using 60 minutes of multi-echo fMRI data to define individual functional networks, however, we also acknowledge several limitations. With 24 adult-child pairs, our sample size is relatively large for precision functional mapping, but small for detecting subtle group differences and thus replication in larger, independent datasets is needed to confirm our findings. While the narrow age range of children in this study (6 – 8 years) can be a strength as it enables focus on a specific developmental period, we cannot make inferences about other developmental periods or longitudinal change. Therefore, future work with longitudinal designs is needed to provide a complete picture of developmental network organization. We also note that network nomenclature remains inconsistent across the field (Uddin et al, 2019, 2023), and although we provide alternative taxonomy for our dataset, this may further impact interpretability of individualized network findings. Though attention and engagement were sustained similarly across groups in our sample (Rai et al., 2025), differences noted in this study between children and adults may be driven by age-related variability in stimulus processing (Campbell et al., 2015; Cantlon & Li, 2013; Hutchison & Morton, 2015). Of note, motion artifacts remain a significant concern in developmental neuroimaging (Frew et al., 2022) and residual motion effects may still influence results seen here, especially given the high-motion children group in our study. Lastly, while new packages such as VertexWiseR (Billaud & Yu, 2024) and FEMA (Parekh et al., 2024) have begun to address the multiple comparisons correction issue in surface-based and vertex wise analyses, currently they lack control in model specification for categorical variables and the ability to mask regions. Hence in this study, both the mixed-effect and logistic regression model report uncorrected p-values, and the observed entropy and assignment differences should be interpreted as trends.

Overall, this work adds to our understanding of age and familial effects on functional topography and connectivity, and our findings highlight the potential for individualized mapping approaches to improve sensitivity of functional connectivity analyses.

## Data and Code availability

Preprocessed and surface projected PreciseKIDS data, descriptions of the viewing conditions used, and scripts created for this study (Python version 3.12.2, MATLAB version 9.11.0 R2021b, R version 4.2.2) are available for download at https://github.com/BrayNeuroimagingLab/BNL_open.

## Acknowledgements

We would like to acknowledge the dedication of all the families that participated in our study, along with the support of the Diagnostic Imaging staff at the Alberta Children’s Hospital.

## Supplementary Material

All supplemental materials are available as a separate document.

