## Supplemental Materials for "Individualized network topography in pre-adolescent children and adults using naturalistic precision fMRI"

**Supplemental material**

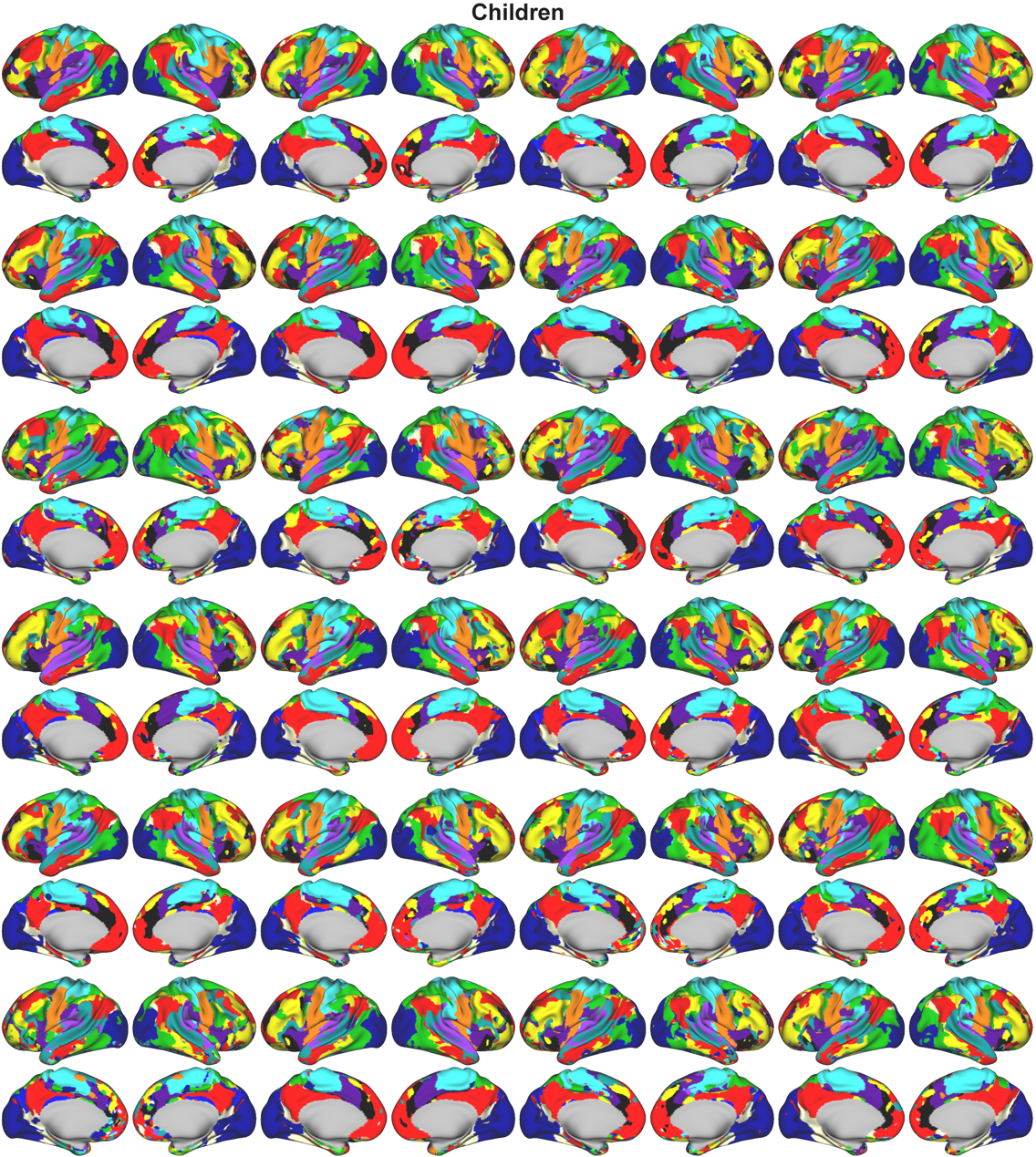

**Supplemental Figure 1: Individualized functional networks for children.** Surface maps are shown for each child within the PreciseKIDS dataset using the template-matching procedure.

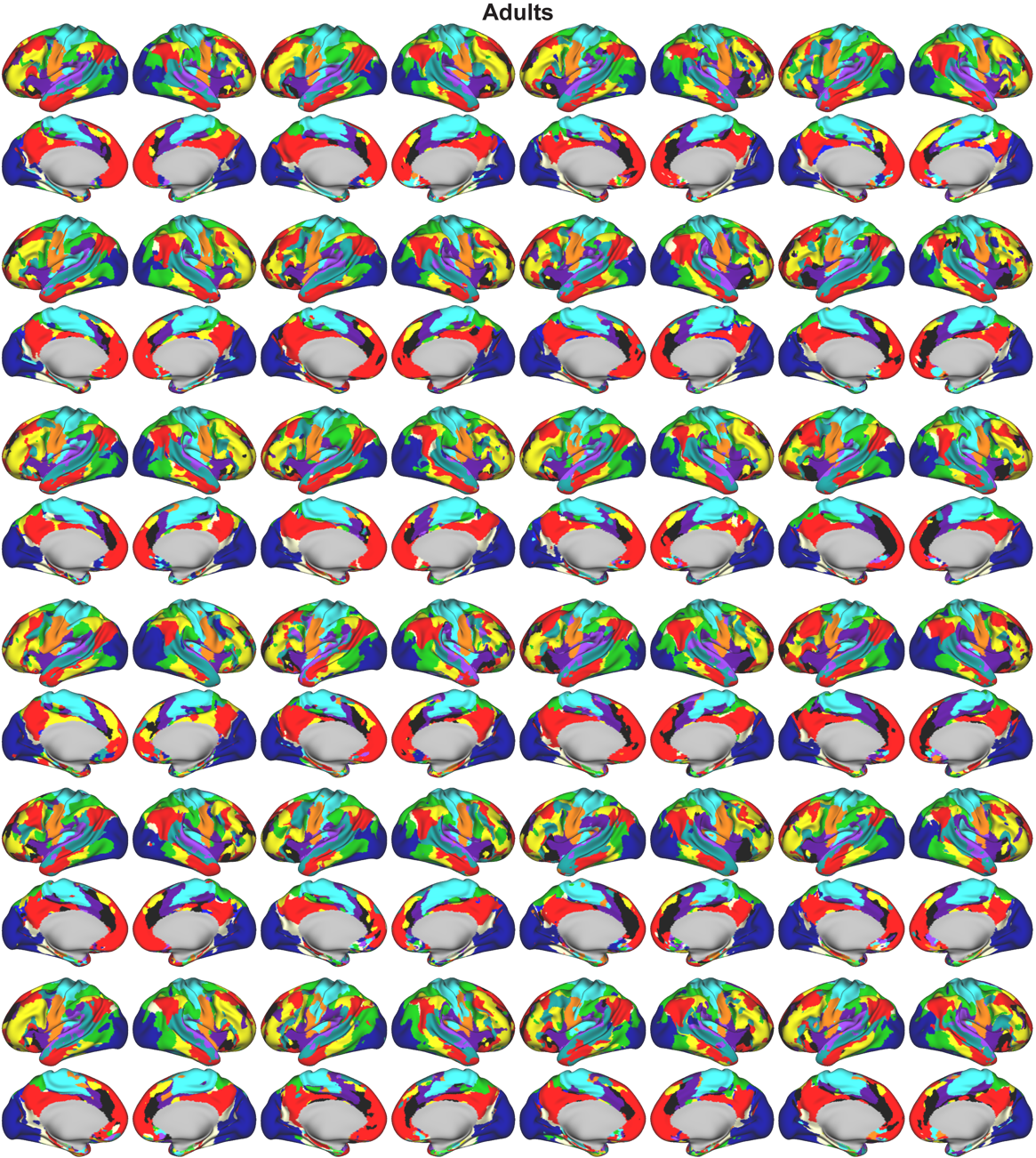

**Supplemental Figure 2: Individualized functional networks for adults.** Surface maps are shown for each adult within the PreciseKIDS dataset using the template-matching procedure.

| **Group** | **Network** | **EG17** | **MG360J12** | **TY17** | **WS90_14** |
| --- | --- | --- | --- | --- | --- |
| Adults | AUD | Auditory (0.55) | Auditory (0.61) | SomatomotorB (0.43) | Auditory (0.26) |
| Children | AUD | Auditory (0.57) | Auditory (0.59) | SomatomotorB (0.44) | Auditory (0.22) |
| Adults | CON | CingOperc (0.54) | CingOperc (0.48) | Sal/VenAttnA (0.52) | AntSal (0.23) |
| Children | CON | CingOperc (0.48) | CingOperc (0.43) | Sal/VenAttnA (0.48) | AntSal (0.24) |
| Adults | DAN | DorsAttn (0.55) | DorsAttn (0.41) | DorsAttnA (0.41) | Visuospatial (0.35) |
| Children | DAN | DorsAttn (0.54) | DorsAttn (0.38) | DorsAttnA (0.41) | Visuospatial (0.32) |
| Adults | DMN | Default (0.66) | Default (0.63) | DefaultA (0.50) | DorsalDMN (0.32) |
| Children | DMN | Default (0.64) | Default (0.59) | DefaultA (0.51) | DorsalDMN (0.31) |
| Adults | FP | FrontPar (0.56) | FrontPar (0.51) | ControlB (0.42) | RECN (0.25) |
| Children | FP | FrontPar (0.54) | FrontPar (0.50) | ControlB (0.40) | RECN (0.22) |
| Adults | MTL | AntMTL (0.13) | VentMulti (0.17) | LimbicA (0.11) | DorsalDMN (0.00) |
| Children | MTL | AntMTL (0.14) | VentMulti (0.21) | LimbicA (0.13) | DorsalDMN (0.00) |
| Adults | PMN | ParMemory (0.16) | FrontPar (0.03) | ControlC (0.11) | Precuneus (0.11) |
| Children | PMN | ParMemory (0.32) | FrontPar (0.07) | ControlC (0.23) | Precuneus (0.25) |
| Adults | PON | Context (0.48) | Default (0.14) | DefaultC (0.51) | VentralDMN (0.24) |
| Children | PON | Context (0.48) | Default (0.16) | DefaultC (0.51) | VentralDMN (0.24) |
| Adults | Sal | Salience (0.46) | CingOperc (0.17) | Sal/VenAttnB (0.30) | AntSal (0.25) |
| Children | Sal | Salience (0.48) | OrbitAffective (0.17) | Sal/VenAttnB (0.28) | AntSal (0.21) |
| Adults | SMd | FootSM (0.49) | Somatomotor (0.73) | SomatomotorA (0.74) | Sensorimotor (0.26) |
| Children | SMd | FootSM (0.50) | Somatomotor (0.67) | SomatomotorA (0.73) | Sensorimotor (0.24) |
| Adults | SML | FaceSM (0.63) | Somatomotor (0.31) | SomatomotorB (0.54) | Sensorimotor (0.15) |
| Children | SML | FaceSM (0.63) | Somatomotor (0.36) | SomatomotorB (0.56) | Sensorimotor (0.20) |
| Adults | Tpole | AntMTL (0.05) | OrbitAffective (0.03) | LimbicA (0.05) | Visuospatial (0.00) |
| Children | Tpole | AntMTL (0.06) | OrbitAffective (0.03) | LimbicA (0.05) | Visuospatial (0.00) |
| Adults | VAN | Language (0.56) | Language (0.47) | TempPar (0.44) | Language (0.30) |
| Children | VAN | Language (0.53) | Language (0.46) | TempPar (0.42) | Language (0.31) |
| Adults | Vis | LatVis (0.70) | Visual2 (0.68) | VisualA (0.60) | HighVisual (0.23) |
| Children | Vis | LatVis (0.68) | Visual2 (0.65) | VisualA (0.59) | HighVisual (0.22) |

**Supplemental Table 1: Alternative network nomenclature using the Network Correspondence Toolbox.** Using the highest Dice matches across four reference atlases, (Glasser et al., 2016; Gordon et al., 2016; Shirer et al., 2012; Yeo et al., 2011), we provide network labels that are commonly used in the field to describe the same cortical networks we have used in this study.

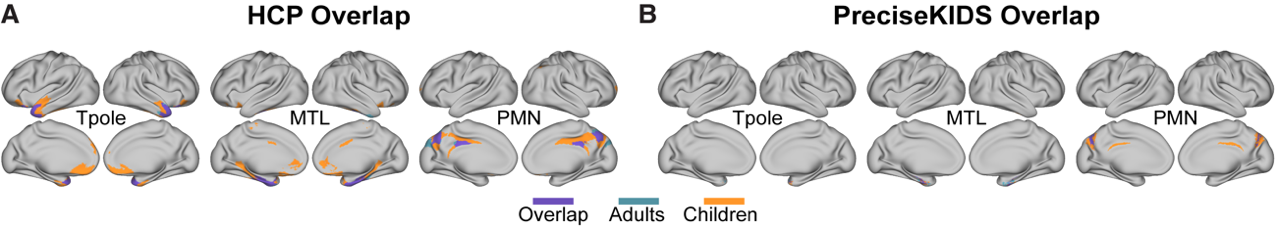

**Supplemental Figure 3: HCP and PreciseKIDS overlap surface maps for smaller networks.** Surface maps are shown across adult and children using both the HCP and PreciseKIDS data for Tpole, MTL, and PMN.

| **Network** | **Children surface area (mm^2^)** | **Adults surface area (mm^2^)** | **Direction** |
| --- | --- | --- | --- |
| DMN | 90468.7 | 86945.3 | Children > Adults |
| VIS | 71664.6 | 62824.3 | Children > Adults |
| FP | 81417.3 | 73545.2 | Children > Adults |
| DAN | 83314.3 | 71722.3 | Children > Adults |
| VAN | 57765.5 | 51133.2 | Children > Adults |
| SAL | 32088.3 | 36188.6 | Adults > Children |
| CON | 50105.1 | 42939.4 | Children > Adults |
| SMd | 58679.3 | 52553.9 | Children > Adults |
| SMl | 35579.9 | 27929.8 | Children > Adults |
| AUD | 28988.1 | 27769.7 | Children > Adults |
| Tpole | 3531.3 | 2323.8 | Children > Adults |
| MTL | 5987.2 | 5346.9 | Children > Adults |
| PMN | 12006.9 | 6938.8 | Children > Adults |
| PON | 27485.1 | 22241.9 | Children > Adults |

**Supplemental Table 2: Cortical surface area differences calculated for any child and any adult in each group across all networks.**

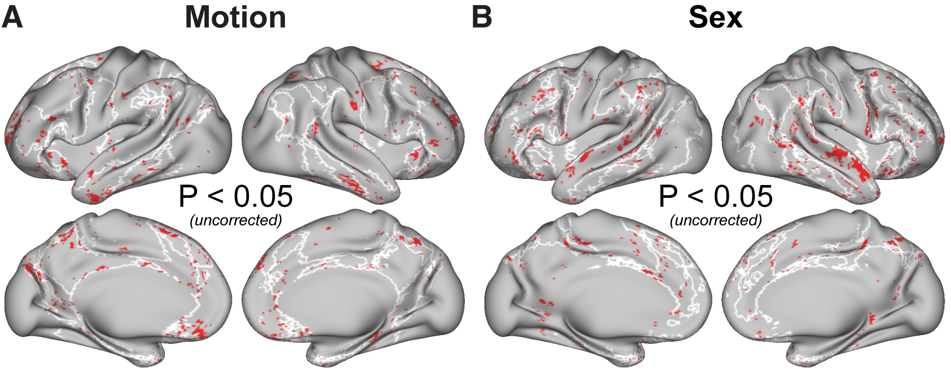

**Supplemental Figure 4: Uncorrected significant effects of motion and sex across children and adults in network assignment.** Using a mixed effect logistic regression model, all vertices with motion (A) and sex (B) effects are shown in red. We found networks with the largest motion effect were DMN, FP, SMd, and PON (p-uncorrected < 0.05) and networks with the largest effect of sex were FP, DAN, VAN, and SAL (p-uncorrected < 0.05), outlined in white in A and B, respectively.

| **Network** | **Term** | **Estimate** | **SD Error** | **Statistic** | **P value** | **Sig.** | **P-adj. value** | **Sig. (FDR)** |
| --- | --- | --- | --- | --- | --- | --- | --- | --- |
| SMl | Group | -7.20E-03 | 3.49E-03 | -2.0629 | 0.0451 | * | 0.247811 |  |
| PON | Group | -6.90E-03 | 2.97E-03 | -2.3209 | 0.0275 | * | 0.247811 |  |
| SAL | Group | -8.12E-03 | 4.56E-03 | -1.7818 | 0.0836 |  | 0.306652 |  |
| DMN | Group | 8.75E-03 | 6.86E-03 | 1.274 | 0.2124 |  | 0.445043 |  |
| VIS | Group | 7.02E-03 | 5.93E-03 | 1.184 | 0.2428 |  | 0.445043 |  |
| VAN | Group | 6.69E-03 | 5.00E-03 | 1.3391 | 0.1874 |  | 0.445043 |  |
| CON | Group | 4.27E-03 | 5.48E-03 | 0.7788 | 0.441 |  | 0.693003 |  |
| FP | Group | -1.99E-03 | 5.91E-03 | -0.3369 | 0.7387 |  | 0.848153 |  |
| DAN | Group | -1.83E-03 | 6.24E-03 | -0.2937 | 0.771 |  | 0.848153 |  |
| SMd | Group | 1.71E-03 | 5.25E-03 | 0.325 | 0.7475 |  | 0.848153 |  |
| AUD | Group | 3.17E-04 | 3.49E-03 | 0.0909 | 0.928 |  | 0.928008 |  |
| FP | Sex (M) | 1.12E-02 | 4.58E-03 | 2.4582 | 0.0197 | * | 0.077332 |  |
| VAN | Sex (M) | 8.91E-03 | 3.73E-03 | 2.3921 | 0.0211 | * | 0.077332 |  |
| AUD | Sex (M) | -7.26E-03 | 2.60E-03 | -2.7912 | 0.0077 | ** | 0.077332 |  |
| DMN | Sex (M) | -4.83E-03 | 5.25E-03 | -0.9197 | 0.3641 |  | 0.667452 |  |
| DAN | Sex (M) | 4.94E-03 | 4.82E-03 | 1.0252 | 0.313 |  | 0.667452 |  |
| CON | Sex (M) | -3.95E-03 | 4.09E-03 | -0.9655 | 0.3399 |  | 0.667452 |  |
| SMl | Sex (M) | -1.78E-03 | 2.60E-03 | -0.6846 | 0.4972 |  | 0.689901 |  |
| PON | Sex (M) | 1.55E-03 | 2.28E-03 | 0.6791 | 0.5017 |  | 0.689901 |  |
| SAL | Sex (M) | -1.93E-03 | 3.44E-03 | -0.5617 | 0.5775 |  | 0.705816 |  |
| SMd | Sex (M) | -1.57E-03 | 4.03E-03 | -0.3889 | 0.6998 |  | 0.769821 |  |
| VIS | Sex (M) | -7.03E-04 | 4.42E-03 | -0.159 | 0.8744 |  | 0.874362 |  |
| SAL | Motion | 1.17E-05 | 3.04E-06 | 3.834 | 0.0004 | *** | 0.004626 | ** |
| DAN | Motion | -8.87E-06 | 4.34E-06 | -2.0465 | 0.0483 | * | 0.265534 |  |
| DMN | Motion | 6.35E-06 | 4.67E-06 | 1.3592 | 0.1821 |  | 0.488193 |  |
| VAN | Motion | -4.31E-06 | 3.27E-06 | -1.3171 | 0.1946 |  | 0.488193 |  |
| SMd | Motion | 4.48E-06 | 3.61E-06 | 1.2427 | 0.2219 |  | 0.488193 |  |
| FP | Motion | -3.77E-06 | 4.12E-06 | -0.9159 | 0.3661 |  | 0.671247 |  |
| AUD | Motion | -1.40E-06 | 2.28E-06 | -0.612 | 0.5437 |  | 0.854407 |  |
| CON | Motion | -1.75E-06 | 3.60E-06 | -0.4865 | 0.6291 |  | 0.864965 |  |
| VIS | Motion | -4.64E-07 | 3.88E-06 | -0.1196 | 0.9053 |  | 0.905316 |  |
| SMl | Motion | -3.11E-07 | 2.28E-06 | -0.1359 | 0.8925 |  | 0.905316 |  |
| PON | Motion | 6.73E-07 | 2.04E-06 | 0.3303 | 0.743 |  | 0.905316 |  |

**Supplemental Table 3: Effects of age group, sex and head motion on surface area differences across networks.**

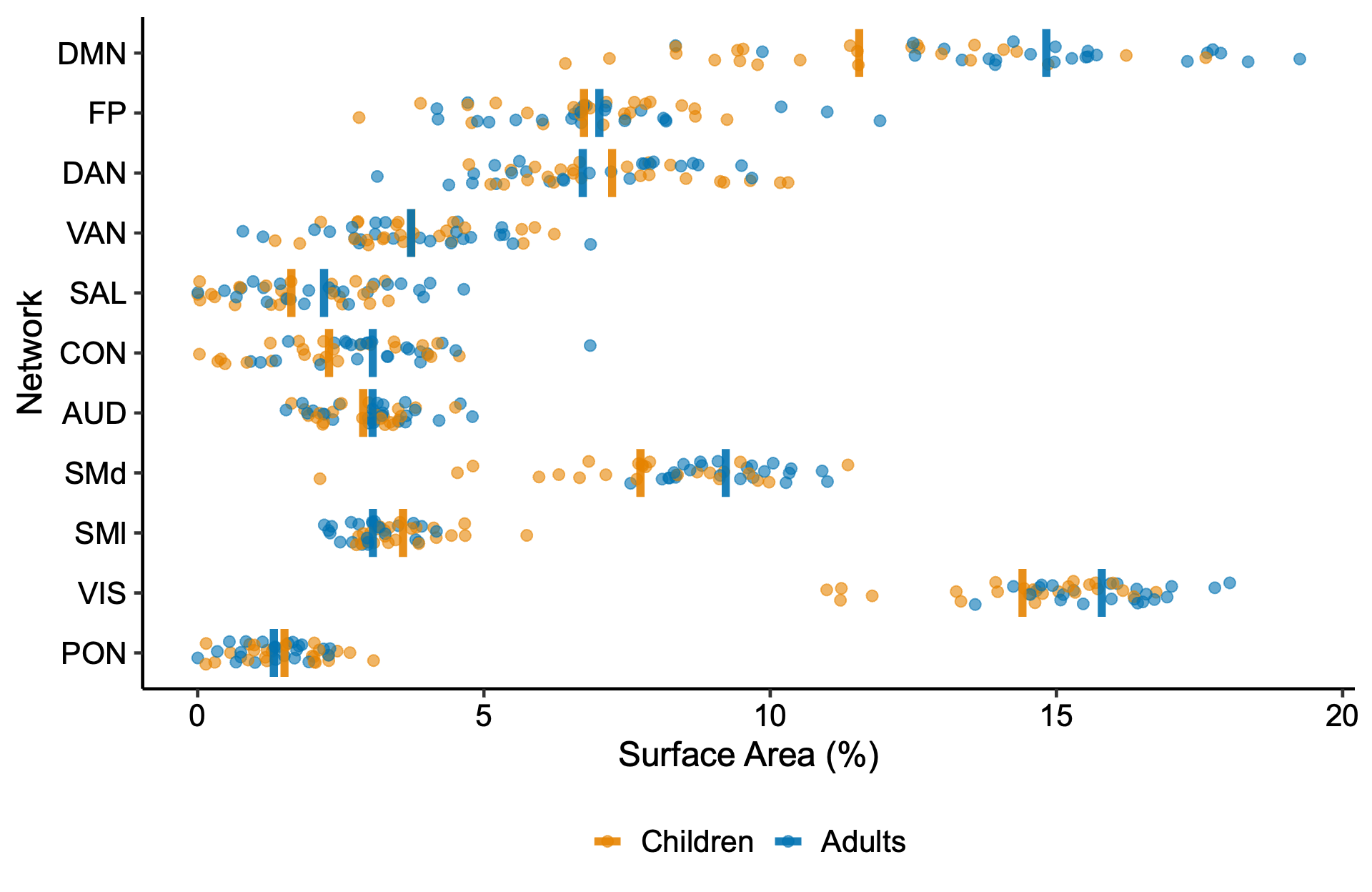

**Supplemental Figure 5: High-confidence surface area comparisons between adults and children across networks.** Surface area differences between high-confidence regions (Dice ≥ 0.3) did not show significant age group effects, however the SAL network showed motion effects (pFDR = 0.009).

| **Network** | **Term** | **Estimate** | **SD Error** | **Statistic** | **P value** | **Sig.** | **P-adj. value** | **Sig. (FDR)** |
| --- | --- | --- | --- | --- | --- | --- | --- | --- |
| DMN | Group | 1.86E-02 | 7.89E-03 | 2.3543 | 0.0256 | * | 0.177896 |  |
| SMd | Group | 1.05E-02 | 4.95E-03 | 2.1231 | 0.0427 | * | 0.177896 |  |
| SMl | Group | -5.20E-03 | 2.62E-03 | -1.9859 | 0.0533 |  | 0.177896 |  |
| PON | Group | -4.70E-03 | 2.45E-03 | -1.9212 | 0.0647 |  | 0.177896 |  |
| CON | Group | 5.55E-03 | 4.57E-03 | 1.2143 | 0.2333 |  | 0.513355 |  |
| VIS | Group | 5.53E-03 | 5.32E-03 | 1.0379 | 0.305 |  | 0.518519 |  |
| DAN | Group | -4.97E-03 | 5.53E-03 | -0.8973 | 0.3771 |  | 0.518519 |  |
| SAL | Group | -3.38E-03 | 3.60E-03 | -0.9384 | 0.3554 |  | 0.518519 |  |
| AUD | Group | 1.69E-03 | 3.00E-03 | 0.562 | 0.577 |  | 0.705188 |  |
| FP | Group | -1.98E-03 | 6.09E-03 | -0.3256 | 0.747 |  | 0.821647 |  |
| VAN | Group | 9.39E-04 | 5.24E-03 | 0.1792 | 0.8587 |  | 0.858738 |  |
| VAN | Sex (M) | 8.06E-03 | 3.91E-03 | 2.0635 | 0.0453 | * | 0.249139 |  |
| AUD | Sex (M) | -5.22E-03 | 2.23E-03 | -2.3353 | 0.0241 | * | 0.249139 |  |
| CON | Sex (M) | -5.98E-03 | 3.50E-03 | -1.711 | 0.0956 |  | 0.350384 |  |
| DMN | Sex (M) | -9.03E-03 | 6.12E-03 | -1.4763 | 0.1499 |  | 0.412179 |  |
| VIS | Sex (M) | -2.97E-03 | 3.97E-03 | -0.7489 | 0.4579 |  | 0.682495 |  |
| FP | Sex (M) | 2.65E-03 | 4.69E-03 | 0.5649 | 0.5758 |  | 0.682495 |  |
| SAL | Sex (M) | -1.96E-03 | 2.77E-03 | -0.7077 | 0.484 |  | 0.682495 |  |
| SMd | Sex (M) | -1.92E-03 | 3.84E-03 | -0.5007 | 0.6202 |  | 0.682495 |  |
| SMl | Sex (M) | -9.73E-04 | 1.95E-03 | -0.4987 | 0.6205 |  | 0.682495 |  |
| PON | Sex (M) | 1.39E-03 | 1.89E-03 | 0.7336 | 0.4685 |  | 0.682495 |  |
| DAN | Sex (M) | 9.03E-04 | 4.29E-03 | 0.2106 | 0.8346 |  | 0.834559 |  |
| SAL | Motion | 9.07E-06 | 2.48E-06 | 3.657 | 0.0008 | *** | 0.008686 | ** |
| DMN | Motion | 1.46E-05 | 5.51E-06 | 2.6428 | 0.0123 | * | 0.067882 |  |
| VIS | Motion | 8.41E-06 | 3.48E-06 | 2.4129 | 0.0201 | * | 0.073547 |  |
| PON | Motion | 2.68E-06 | 1.70E-06 | 1.5815 | 0.1227 |  | 0.337504 |  |
| SMd | Motion | 4.47E-06 | 3.47E-06 | 1.2867 | 0.2071 |  | 0.455686 |  |
| FP | Motion | 4.36E-06 | 4.21E-06 | 1.0359 | 0.307 |  | 0.562821 |  |
| CON | Motion | 2.52E-06 | 3.12E-06 | 0.8079 | 0.424 |  | 0.666337 |  |
| VAN | Motion | -1.62E-06 | 3.43E-06 | -0.4708 | 0.6402 |  | 0.880252 |  |
| DAN | Motion | -2.51E-07 | 3.86E-06 | -0.0649 | 0.9486 |  | 0.993218 |  |
| SMl | Motion | 1.47E-08 | 1.71E-06 | 0.0085 | 0.9932 |  | 0.993218 |  |
| AUD | Motion | 3.76E-07 | 1.96E-06 | 0.1915 | 0.849 |  | 0.993218 |  |

**Supplemental Table 4: High-confidence surface area significant differences between adults and children across networks.**

| **Contrast** | **Network** | **Estimate** | **SD Error** | **Z ratio** | **P value** | **Sig.** |
| --- | --- | --- | --- | --- | --- | --- |
| LMA-LMA – LMC-LMC | DMN | 0.02226 | 0.01429 | 1.55709 | 0.11945 |  |
| LMA-LMA – LMC-LMC | VIS | 0.01672 | 0.01429 | 1.16985 | 0.24206 |  |
| LMA-LMA – LMC-LMC | FP | -0.00101 | 0.01429 | -0.0704 | 0.94386 |  |
| LMA-LMA – LMC-LMC | DAN | 0.04893 | 0.01429 | 3.42316 | 0.00062 | *** |
| LMA-LMA – LMC-LMC | VAN | 0.01664 | 0.01429 | 1.16398 | 0.24443 |  |
| LMA-LMA – LMC-LMC | SAL | -0.10484 | 0.01429 | -7.3339 | 0.00000 | *** |
| LMA-LMA – LMC-LMC | CON | 0.07126 | 0.01429 | 4.98496 | 0.00000 | *** |
| LMA-LMA – LMC-LMC | SMd | 0.08286 | 0.01429 | 5.79630 | 0.00000 | *** |
| LMA-LMA – LMC-LMC | SMl | 0.00862 | 0.01429 | 0.60269 | 0.54671 |  |
| LMA-LMA – LMC-LMC | AUD | -0.00834 | 0.01429 | -0.5836 | 0.55946 |  |
| LMA-LMA – LMC-LMC | PON | 0.00477 | 0.01429 | 0.33369 | 0.73862 |  |

**Supplemental Table 5: *Post-hoc* comparisons of network topography similarity across low-motion adults and low-motion children.**

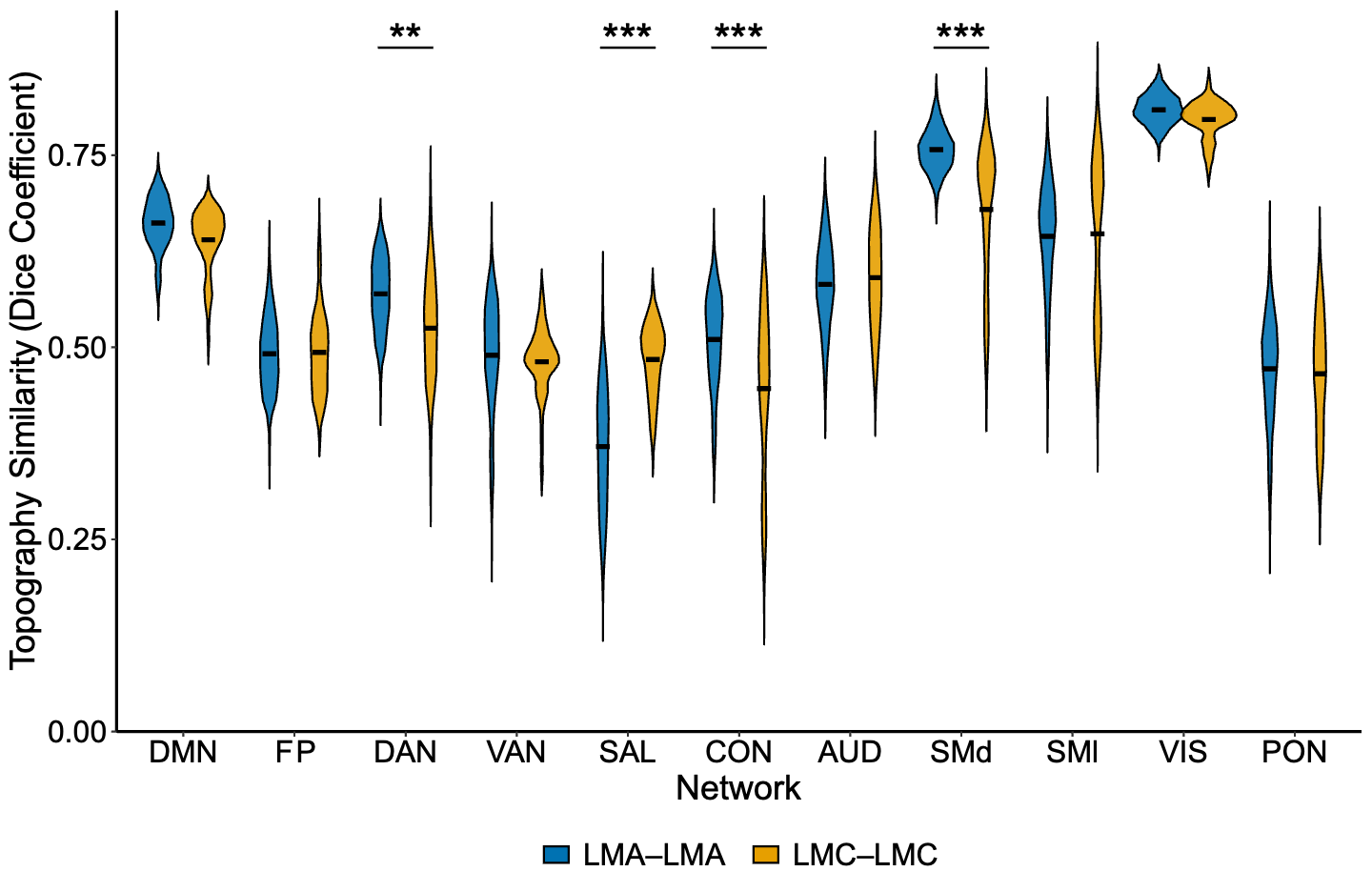

**Supplemental Figure 6: Within-group similarity of network topography in reliability matched low-motion adults and low-motion children.** Pairwise dice similarity in network assignment for low-motion adults (LMA-LMA; blue) with 30 minutes of post-censored data and low-motion children (LMC-LMC; orange) with 60 minutes of data for each network. There was a significant effect of network (p < 0.002) and interaction between network and group (p < 0.02). Asterisks denote *post-hoc* comparisons with FDR correction: ** p < 0.01, *** p < 0.001.

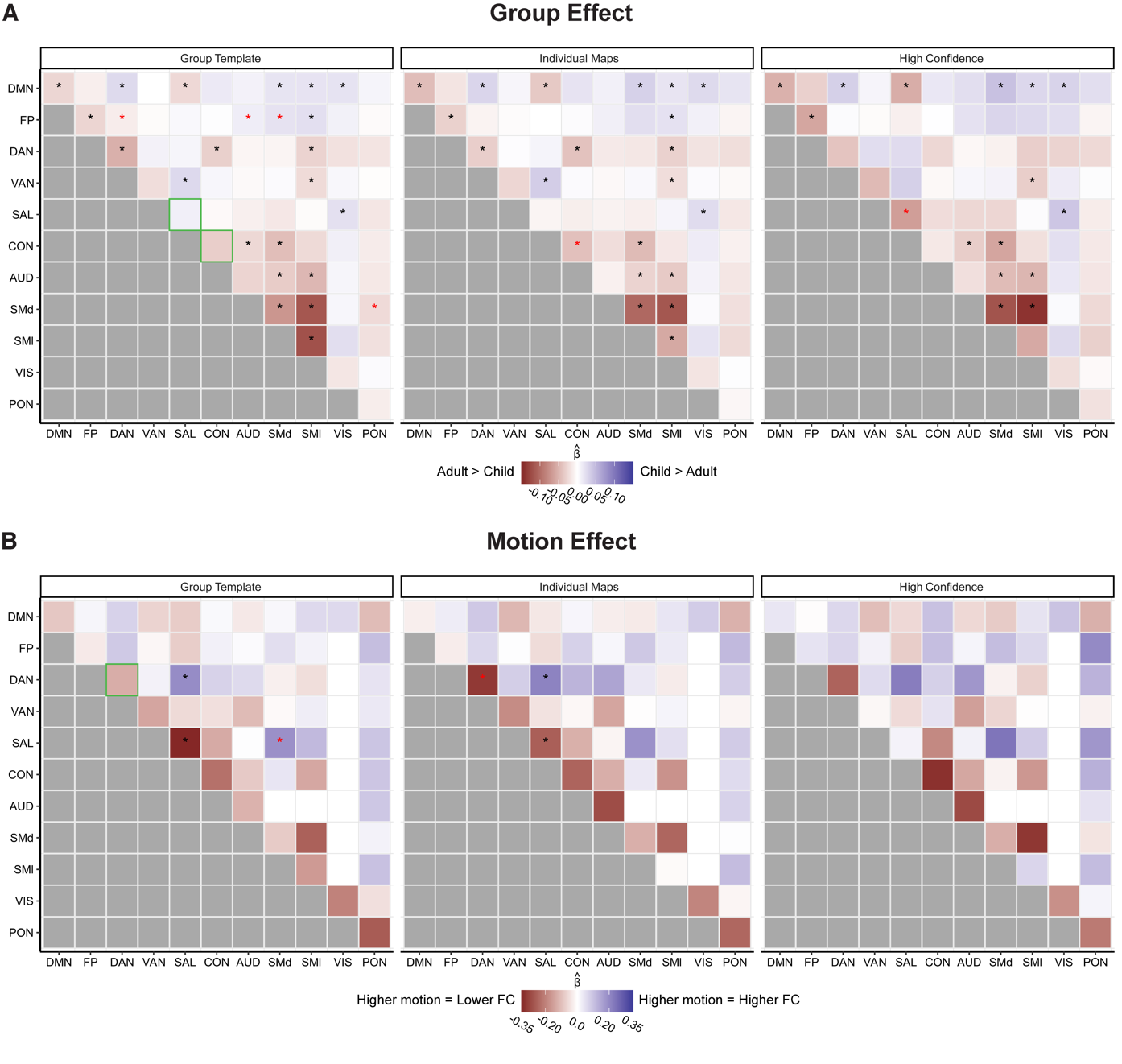

**Supplemental Figure 7: FC differences between children and adults across networks and mapping approaches using 15-minutes of independent data.** Group templates (left), individualized maps (middle), and high-confidence individualized (Dice ≥ 0.3) (right) approaches were used. A) In red, beta coefficients from the group effect show stronger FC in adults compared to children, and in blue stronger FC in children is shown compared to adults. Across all approaches, adults had greater within-network FC than children. B) Motion effects in red indicate higher motion decreased FC estimates (negative betas), and effects in blue indicate motion increased FC estimates (positive betas). Significance asterisks denote comparisons with FDR correction (p < 0.05). Red stars represent significance for a specific approach only. Green boxes reflect differences in within-FC between the group template and either of the individualized approaches.
